# Analyzing Genomic Data Using Tensor-Based Orthogonal Polynomials with Application to Synthetic RNAs

**DOI:** 10.1101/2020.04.24.059279

**Authors:** Saba Nafees, Sean H. Rice, Catherine A. Wakeman

## Abstract

An important goal in molecular biology is to quantify both the patterns across a genomic sequence and the relationship between phenotype and underlying sequence. We propose a multivariate tensor-based orthogonal polynomial approach to characterize nucleotides or amino acids in a given sequence and map corresponding phenotypes onto the sequence space. We have applied this method to a previously published case of small transcription activating RNAs (STARs). Covariance patterns along the sequence showcased strong correlations between nucleotides at the ends of the sequence. However, when the phenotype is projected onto the sequence space, this pattern doesn’t emerge. When doing second order analysis and quantifying the functional relationship between the phenotype and pairs of sites along the sequence, we identified sites with high regressions spread across the sequence, indicating potential intramolecular binding. In addition to quantifying interactions between different parts of a sequence, the method quantifies sequence-phenotype interactions at first and higher order levels. We discuss the strengths and constraints of the method and compare it to computational methods such as machine learning approaches. An accompanying command line tool to compute these polynomials is provided. We show proof of concept of this approach and demonstrate its potential application to other biological systems.

## INTRODUCTION

Due to the inherent complexity in biological systems, much of the analysis that aims to study systems, such as sequence structure and function in genomics, has been largely based on computational methods. Advancements in sequencing technology and the rise in the availability of genomic data has led to the development of novel computational tools that seek to determine the relationship between underlying sequence and the resulting function or phenotype. These methods aim to predict protein function given sequence and structure information, identify novel and potential DNA binding motifs, including transcription factor binding sites, and determine RNA secondary structure based on the underlying sequence (1)-(4). While computational methods, such as machine learning models, provide powerful ways to extract patterns in complex data, they lack the clarity of interpretation that comes with purely mathematical approaches. In this work, we present a method to do analytical mathematics with sequences that integrates well with computational methods and that has the potential to reveal new results and complement those given by purely computational or machine learning approaches.

Current methods that aim to connect sequence information to output phenotype largely utilize machine learning tools, such as deep learning, to quantify the role of each site in a sequence in the resulting phenotypes. Some examples of this include deep learning tools that aim to predict protein structure given the underlying amino acid sequence and others that are designed to predict the effect of each site in a pre-mRNA sequence on mRNA splicing (5, 6). Although these methods seem promising and do reveal substantial biological insights, they come with caveats such as the requirement of large training datasets and “the incorporation of irrelevant features” into the deep learning model as in the case of SpliceAI (6).

Here, we describe a mathematical method using multivariate tensor-based orthogonal polynomials to convert sequence information into vectors and build orthogonal polynomials with the aim of quantifying the effect of the sequence states on the resulting phenotype (7). Our method proposes an entirely novel way to work with nucleotides in a genomic sequence. Elements in a sequence (nucleotides, amino acids, *etc*.) are not meaningfully described by scalar values (*i*.*e*. by a single number) or by binary values because they are distinct types of things that are not well-characterized in this way (7).

In general, instead of representing nucleotides with numbers (real or binary), current methods utilize kmers, or words of length k, to represent a set of nucleotide bases. Here, a 1-mer is the same as a single site in a sequence. The use of kmers is ubiquitous as a bioinformatics tool and they’re used widely in many bioinformatics applications such as genome assembly and compressing genomic sequence data (8, 9). In addition, kmers can be utilized to look for associations between different sequences and any associated traits in GWAS studies (10). Another widely used representation is encoding each nucleotide as a one-hot encoded array that is fed into machine learning algorithms. One-hot encoding refers to denoting a given nucleotide at a site with a 1 and the rest of the nucleotides as zeros. Many of these algorithms are deep learning methods that are based on neural networks with many layers that attempt to understand the underlying variation in sequence data and predict phenotypes (11).

In our system, we represent monomers not with individual numbers or sets of words (kmers), but with vectors. This is similar to one-hot encoding of nucleotides, in which a nucleotide is represented as an array to be used in computational genomics or machine learning methods. In our approach, however, the array is treated mathematically as a vector (a tensor of rank one). Note that a vector here represents a single site, not a sequence; a pair of sites is captured with a tensor of rank 2 (a matrix). These vectors and matrices can then be incorporated into mathematical functions, such as polynomials. In contrast to computational approaches, denoting nucleotides as mathematical objects in this way allows us to do formal mathematics with sequence information. For example, the mean vector at a site gives the distribution of nucleotides at that site, same as what a sequence logo plot gives, and the mean outer product of two sites gives us a matrix that is the covariance matrix (see Methods). After denoting each site in a sequence as a vector (instead of the entire sequence being a vector), we then build orthogonal polynomials based on these vectors and project the phenotypes onto the polynomial space. This reveals not only first order interactions between parts of sequences but also quantifies higher order effects of having particular nucleotides at given sites. Thus, doing purely mathematical analyses with this representation of nucleotides yields biologically meaningful results. Note that both the machine learning and the mathematical approaches share the common fundamental step of one-hot encoding the nucleotides but this is in contrast to kmer methods which denote sets of nucleotides as words of length k.

In the case of RNA sequence analysis, secondary structure prediction is an active area of development. For example, predicting pseudoknots without constraints is impossible and is an NP-complete problem (12). RNAs are extremely versatile molecules and play important regulatory roles in gene expression by affecting transcription and translation through various different mechanisms, including intermolecular (trans) and intramolecular (cis) interactions (13). Of particular importance is the ability of RNA to control transcription initiation or termination through secondary structure formation of hairpins and loops (14). In recent years, much work has been done to understand the role of sequence composition in the regulatory potential of RNAs by not only studying the ones that occur naturally in bacterial systems but also synthetically constructing these regulators de novo (15, 16). This presents a unique opportunity to employ novel mathematical tools to quantify the sequence-function relationship between RNAs and their resulting regulatory activity in both synthetic and natural systems.

In general, given sequence information and corresponding phenotype data, one important objective is to quantify exactly how the underlying sequence, whether DNA, RNA or protein, gives rise to variation in phenotype. We use the term “phenotype” broadly, to refer to any measurable biological trait that is (potentially) influenced by one or more sequences. This could include whole organism properties as well as molecular traits, such as the rate of expression of a gene product (as in (17)) or binding propensity of a transcription factor. Though we will use molecular traits as examples – since there is more data available relating these to sequence variation – the methods that we present could be applied in exactly the same way to whole organism traits such as morphology or expression of a disease. In the case of RNA regulators, one measurable phenotype is the regulatory activity of an RNA as captured experimentally through the use of fluorescent reporters (18).

We have applied this method to a case of regulatory RNAs called Small Transcription Activating RNAs (STARs) that were synthetically constructed and whose regulatory activity was quantified experimentally by monitoring levels of green fluorescent protein expression (Figure 1,(19)). Our methods showcase important characteristics about the sequence composition of these synthetic RNAs and the corresponding target RNAs that they bind to. When analyzing just the variation in the given sequences, covariation patterns emerge that indicate interactions between the two ends of the STAR linear region. However, somewhat surprisingly, this pattern does not emerge when the phenotype values (fluorescence indicating transcription termination activity) are projected onto the sequence space. When looking at second order effects and projecting the phenotype onto pairs of nucleotides, we find a few notable interactions spread out along the sequence indicating potential intramolecular binding. By showing proof of concept of this method applied to these synthetic RNAs, we demonstrate its ability to capture nucleotide (or amino acids in the case of proteins) interactions across sequences and quantify sequence-phenotype interactions can be leveraged when applied to other biological systems.

**Figure 1.**
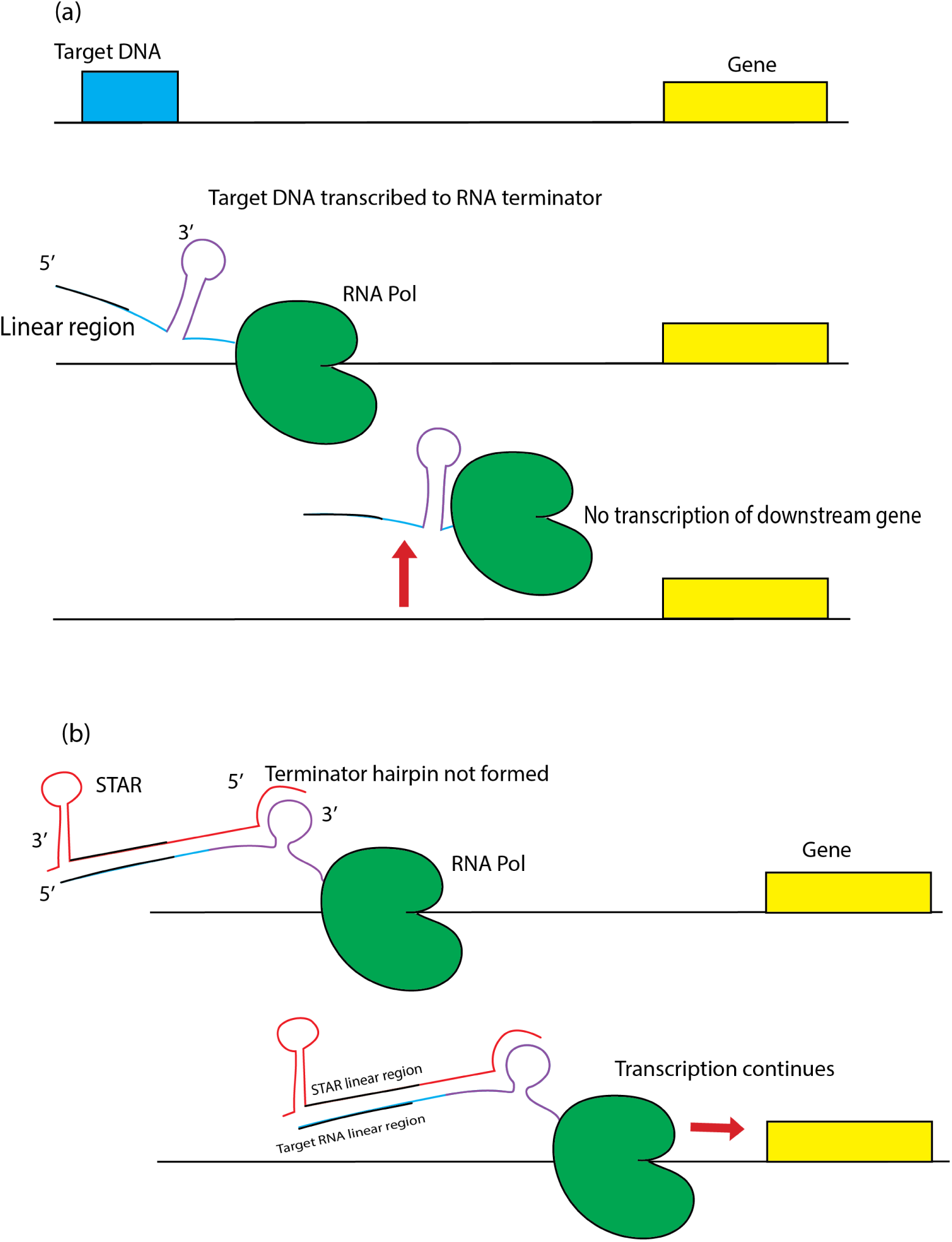
STAR and Target RNA mechanism. (a) illustrates the case in which a target DNA sequence is transcribed into an intrinsic terminator hairpin which displaces the polymerase, preventing transcription of the downstream gene. (b) shows that when STAR binds to the target RNA, the terminator hairpin is prevented from forming and transcription of the downstream gene continues. The purple region in the target RNA indicates the terminator helix which is disrupted upon binding of STAR. The 40 nucleotide linear region in both the target RNA and STAR is depicted with a black line. This was the sequence that was varied while the terminator hairpin sequence remained the same across all 99 variants. Figure adapted from (19).

The case of STAR RNAs is particularly interesting to study at the sequence level due to the impact that both intra- and intermolecular interactions may have on the function of this type of RNA. Our method described herein enables prediction of nucleotide sequence covariation both positively and negatively impacting the function of this RNA-based regulator. These findings have the potential to uncover critical intra- and intermolecular interactions within the STAR RNA regulatory system. The impact of nucleotide covariation on both intra- and intermolecular interactions is highlighted in the examples from Figure 2 depicting the ability of different hairpin sequences to potentially form an intermolecular interaction with an unchanging target sequence. In the scenario shown in Figure 2(a), a favorable intramolecular interaction forming a strong hairpin is likely to prevent the formation of the less favorable intermolecular interaction. In the scenario shown in Figure 2(b), a slight sequence change in the hairpin RNA has shifted the balance to a far more favorable intermolecular interaction likely to outperform the relatively weak intramolecular interaction. Interestingly, a covariation on the other end of the potential hairpin restores a favorable intramolecular interaction without impacting the theoretical strength of the intermolecular interaction (Figure 2c). Despite the theoretical strength of the intermolecular interaction depicted in Figure 2(c), the competing intramolecular structure is likely to impair full intermolecular function. Thus, the study of both the positive and negative impacts of covariation on molecular function can provide insight into the nucleotides involved in important intra- and intermolecular interactions. Importantly, while the Watson-Crick base pairing interactions depicted in Figure 2 are rather easy to computationally predict, there are many other types of interactions occurring within structured RNAs that are less simple to predict. Studies providing insight into the functional importance of sequence covariation may assist in improving the predictions of these other types of structural elements in the future.

**Figure 2.**
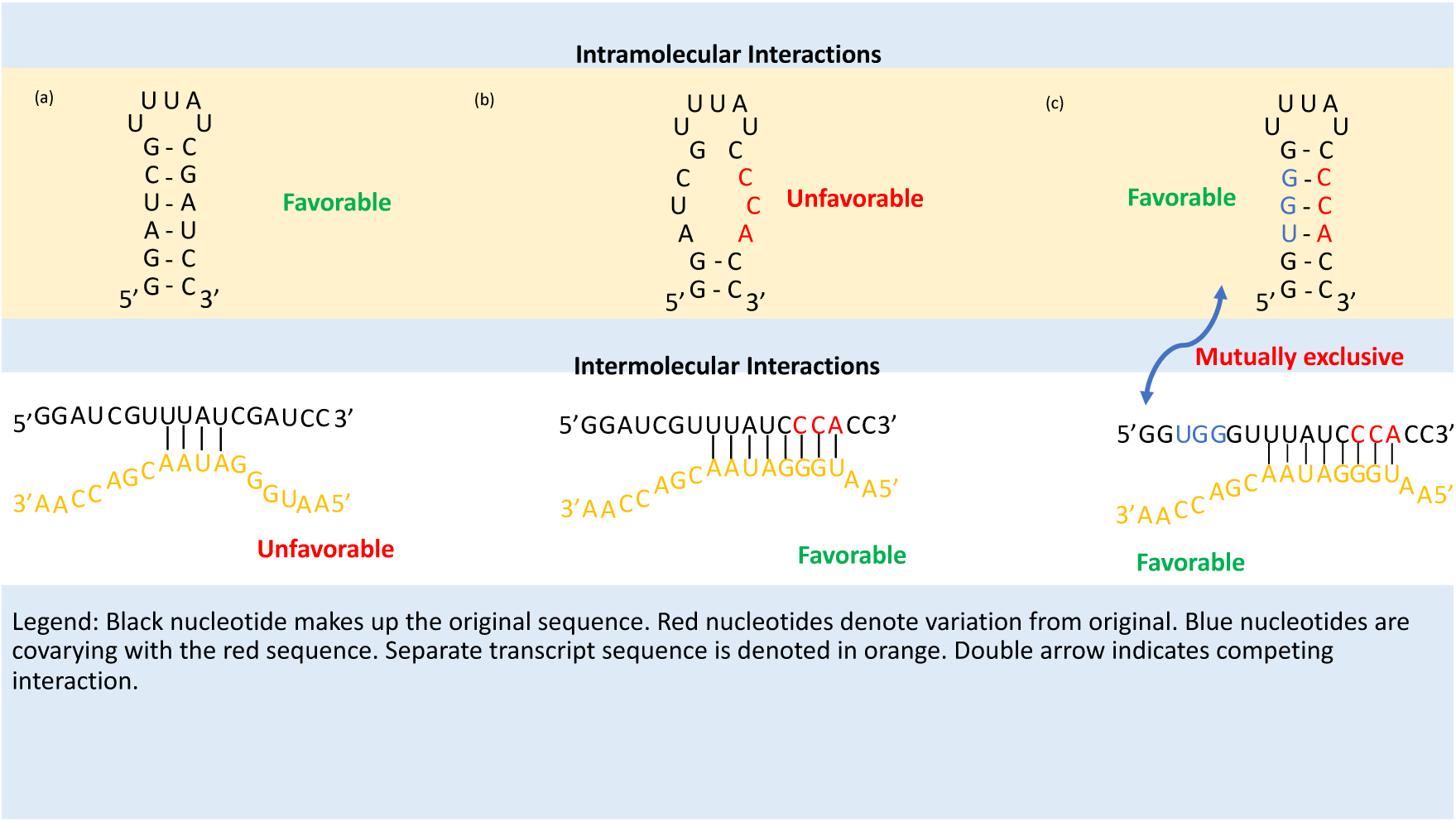
Importance of covariation for secondary structure. (a) shows a case where there exists favorable intramolecular interaction (top panel) and an unfavorable intermolecular interaction with another transcript (bottom panel). (b) Here, the same original sequence has three nucleotides mutated which gives rise to an unfavorable intramolecular interaction (top panel) but a favorable intermolecular interaction with the other transcript (bottom panel). (c) shows mutually exclusive structures with competing intramolecular and intermolecular interactions when there is a mutation (denoted in blue) that covaries with the nucleotides denoted in red.

## MATERIALS AND METHODS

### Application of orthogonal polynomials

Given a set of DNA, RNA, or protein sequences, along with corresponding phenotypic data, our method consists of building tensor-valued first and higher order polynomials and projecting the phenotypic data into this polynomial space (see supplementary methods for detailed explanation and proofs). Note that we assume that the sequences we are working with are of good sequencing quality and depth. Thus, before applying this method to sequence data, the user will have to perform general bioinformatic pre-processing steps to produce the final set of sequences to be analyzed. To apply our methods to the STAR system, we first converted each site in each sequence of the 99 sequences of STAR and target RNAs into a vector as depicted in Figure 3. This is similar to one-hot encoding an array which is a first step when feeding sequence data to a machine learning algorithm (11). For each type of RNA, each sequence had a corresponding experimentally derived OFF/ON fluorescence value which served as the numerical phenotype that we later project onto the polynomial space.

**Figure 3.**
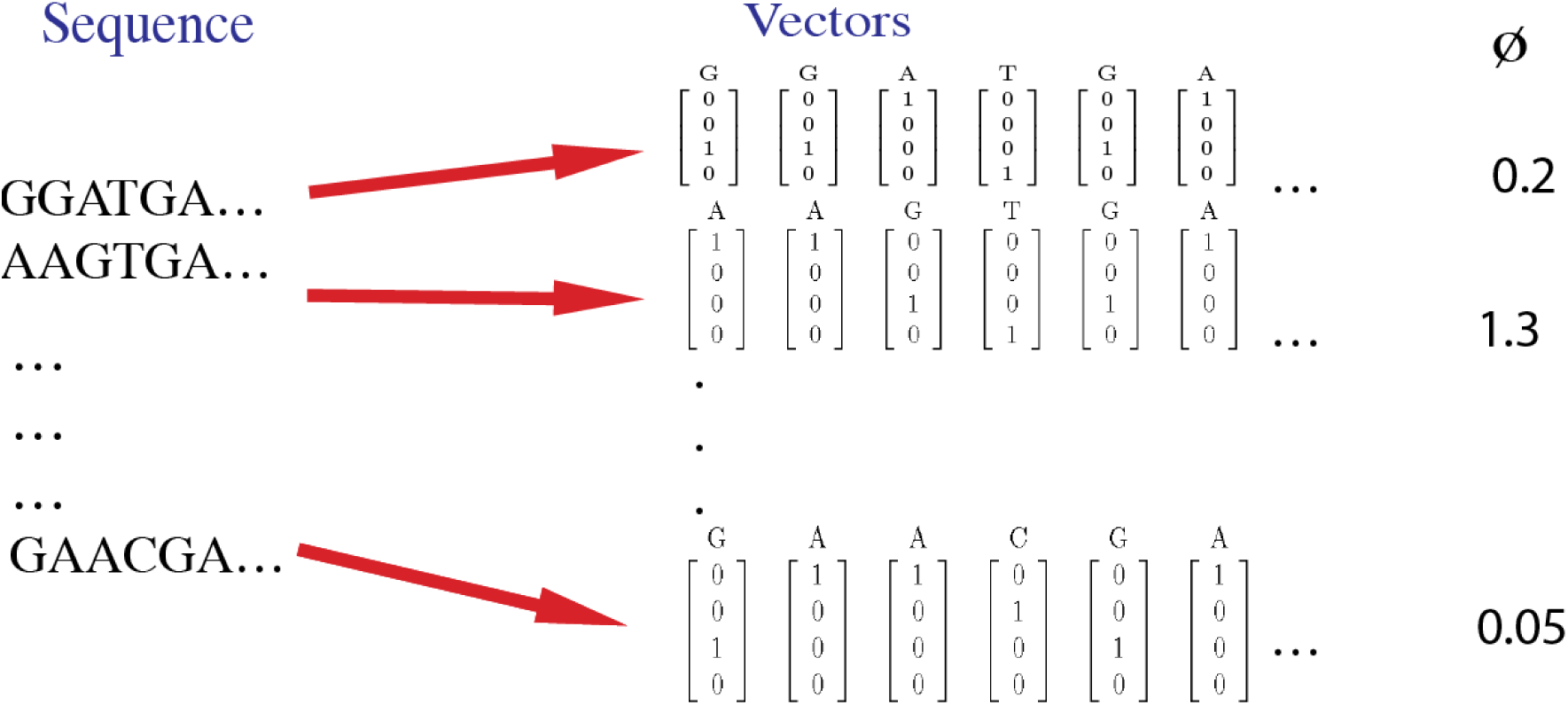
Example sequences with corresponding phenotypic values. Example sequences with corresponding phenotypic values. The first step in our methods is to convert each site in a DNA or RNA sequence to its respective 4-dimensional vector. In this example, a set of 6 sequences is shown, each corresponding to a phenotype (a real-valued number). For DNA and RNA, the vectors are 4-dimensional but this can be changed to a 20-dimensional vector in the case of proteins. The phenotype (*ϕ*) is the off/on fluorescence value associated with the sequence.

After subtracting out the means across all sites in the set of sequences, we get the first order M vectors (see example in described in supplementary methods). We use these to find the variances at each site and covariances between each pair of sites. The covariance analysis shows positive and negative relationships between a pair of two sites. It picks out not only the correlations between sites across a sequence but also the relationship between the *state* at one site (what nucleotide is present) and the state at another site (Figure 3).

The covariance matrix for two sites is the mean, across all individuals, of the outer product of 𝕄^1^ and 𝕄^2^, where 𝕄^1^ and 𝕄^2^ are first order vectors for each individual in the population:

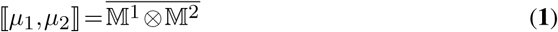

Next, we constructed orthogonal polynomials based on our vectors and projected our variable of interest (OFF/ON values) onto the polynomial space as shown below and detailed in Supplementary Methods.

For scalar traits (body mass, degree of altruism, etc.), we can write some other variable (fitness, susceptibility to a disease, protein folding, *etc*.) as a function of the trait by constructing a set of orthogonal polynomials, of increasing order, for the trait. Orthogonality, here, is defined with respect to the distribution of variation in the population (*i*.*e*. the population distribution is the weight function). This is essentially generalized Fourier analysis.

For example, to write some variable *F* as a function of a single phenotypic trait (*ϕ*), we write the series:

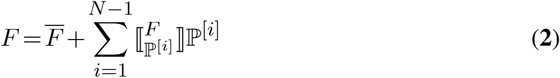

where ℙ^[*i*]^ is the *i*^*th*^ order orthogonal polynomial in *ϕ* (*i*.*e*. it has leading term *ϕ*^*i*^), 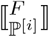 is the projection (regression) of *F* onto polynomial ℙ^[*i*]^, and *N* is population size.

The projection of a variable *F* onto a scalar based polynomial ℙ^*i*^ is just the regression of *F* on ℙ^*i*^ (denoted 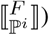) multiplied by ℙ^*i*^. For a vector based polynomial, 𝕄^*i*^, the projection of *F* for a particular individual is given by:

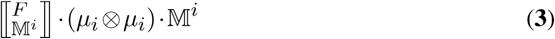

Where “·” represents inner product and “⊗” represents outer product. The role of the new term, *µ*_*i*_ ⊗ *µ*_*i*_, is developed in the section **“Deriving the vector valued equation from the scalar case”** in Supplementary Methods.

For the first order analysis, we are interested in the regression of the phenotype (OFF/ON values) onto the first order conditional polynomial. This is presented here as 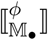 and distributions of these regressions onto target RNA and STAR linear regions are shown in Figures 6 and 7 respectively.

First order analysis is useful to get an overall idea of how a given trait is related to the underlying sequence. However, biological systems are complex and relationships between sequence and the corresponding phenotypic traits often exhibit higher order associations. The challenge with trying to capture higher order relationships, for example, quantifying the regression of the trait on the combination of two or more sites at once, is that this becomes computationally difficult for sequences with large numbers of sites. Thus, second and third order polynomials were constructed for a smaller set of sites. We identified these sites as likely candidates after doing first order analysis and noticing sites that potentially exhibited second and third order interactions. Polynomials up to third order were built for three interacting sites and the regressions of the phenotype on the third orthogonal polynomial were computed (Figures S5-S6). In addition, regressions of the phenotype on the second order orthogonal polynomial were built for six interacting sites in the 5 prime region of the STAR sequence (Figures S7-S10).

All code to construct up to third order orthogonal polynomials is written in the Python programming language. All computational analysis and visualization of results was also done in Python. A command line tool to compute these polynomials based on sequence data and corresponding phenotype data has been constructed and is under further development. Users can clone the Github repository, plug in their own data, and utilize the tool as shown in the repo. To start, users input their sequence data and corresponding phenotype data in separate text files. The program accesses the sequence data (complexity O(1)), converts each letter to a 4-dimensional array, and stores this array as phi (see Supplementary Methods). All the rest of the mathematics is done on these vectors. Supplementary Table 3 shows examples of the computational time and memory requirements to build up to third orthogonal polynomials for STAR/target RNA sequences. It also contains an example of runtimes for a set of 10,000 DNA sequences that are two sites each (see github for more info). In general, the command line interface will run efficiently for shorter sequences compared to longer ones. For instance, when doing just first order analysis for the 40 site STAR sequences, the runtime is efficient but doing second order on all 40 sites is not computationally feasible. One can, however, do second order on sections of the sequence (e.g., 10-site regions) or two sites at a time which is what we have done in this work. The analysis for this paper, along with data and files that generate the figures, has been uploaded to this github project page.

### Application to STARs (small transcription activating RNAs)

We applied our methods to the case of a synthetic RNA regulator designed by (19) known as STARs or small transcription activating RNAs. In this system, a target DNA sequence, containing the necessary information for termination, is placed upstream of a gene. When transcribed, this sequence turns into an intrinsic terminator with a linear region and a hairpin structure. Upon formation of this structure, the polymerase gets knocked off, preventing the transcription of the downstream gene (Figure 1(a), (19, 20)). This is known as the “OFF state”. In the “ON state”, a STAR is constructed such that upon binding to the target RNA, terminator hairpin formation is prevented and the polymerase continues transcription of the downstream gene (Figure 1(b)).

In this system, it was determined that the linear region of the STAR binding to the corresponding part on the target RNA was critical to the activation of the downstream gene. To establish this, linear regions of 100 STAR:target RNA variants were constructed *de novo* using a software package known as NUPACK (21). In this construction, only the linear recognition region was varied while the terminator hairpin remained the same for all variants. This linear region was 40 nucleotides long, as determined experimentally, and it was hypothesized that variation in this part of the sequence would give rise to distinct OFF and ON states for all STAR:target variants (see methods in (19) for details). For the application of our methods to this system, this set of 40 nucleotide long sequences in the STAR and target RNA were used to build first and higher order orthogonal polynomials. Fold and off values were then projected onto the space of the orthogonal polynomials.

## RESULTS

### Covariances showcase nucleotide interactions across STAR sites

As part of first order analysis, means, variances and covariances were computed for all 40 sites across the population of STAR and target RNA sequences. The 40 site long linear region was determined experimentally as being the optimal length to activate transcription (see Methods in (19)). Figure 4 shows frequencies of nucleotides along the 99 sequences of the target RNA. consist of nucleotides cytosine (C), thymine (T), and uracil (U, in the case of RNA). Purines consist of adenine and guanine (G).

**Figure 4.**
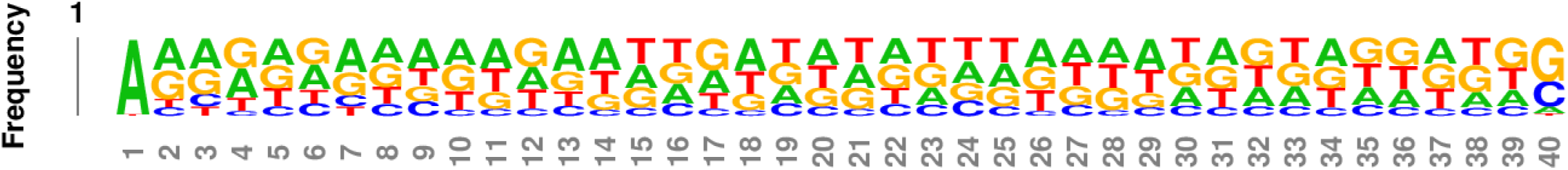
Means of nucleotides present at each site. Frequencies of nucleotides present at each site of the target RNA (from 3 prime to 5 prime) sequence generated by the NUPACK software.

The covariance analysis yields matrices for all pairs of sites, across the 40 sites, and quantifies the relation between having a particular nucleotide at one site and another nucleotide at another site. Since there are 40 sites in this system, there are 780 unique pairs of sites. The covariance between each pair of sites is a matrix and thus, there are 780 4×4 matrices which account for all sites and all possible nucleotides at each site. To understand the distribution of these covariances, a histogram was constructed and different cutoffs were analyzed (Figures S1, S3-S4). Supplementary Figure 1 shows that as the cutoff became larger, the number of “highly covarying” pairs became smaller. The cutoff of −0.05 and 0.05 was selected after testing multiple different cutoffs which showed the same patterns (Figures S3 and S4 showing cutoffs of 0.04 and 0.03 respectively). The cutoff of 0.05 yields 32 site pairs, indicating nucleotides at specific sites covarying highly and positively with each other and nucleotides at other sites covarying highly and negatively with each other (see Supplementary Table 1 for actual values).

These large positive and negative covariances are visualized as seen in Figure 5 and Figure S2 The start of the sequence, depicted as site 1, is the 5’ end of the STAR while the end of the sequence is the 3’ end. It can be immediately noted that sites near the 5’ end of the sequence (sites 2-6) covary with themselves and with all other sites, no other sites covary with anything else except with one of those. And the sites that sites 2-6 connect with are fairly evenly distributed throughout the rest of the sequence. Additionally, there is a lack of notable correlation between sites in the middle portion of the sequence. One possible explanation for this result is that this interaction between sites at the ends of the sequence is preventing any potential binding between the linear region and the hairpin part of the RNA so that the hairpin region can stay intact and pursue its function of terminating transcription of the downstream gene. This type of function for transacting regulatory RNAs has been well described in other contexts ((19),(22) - (24)).

**Figure 5.**
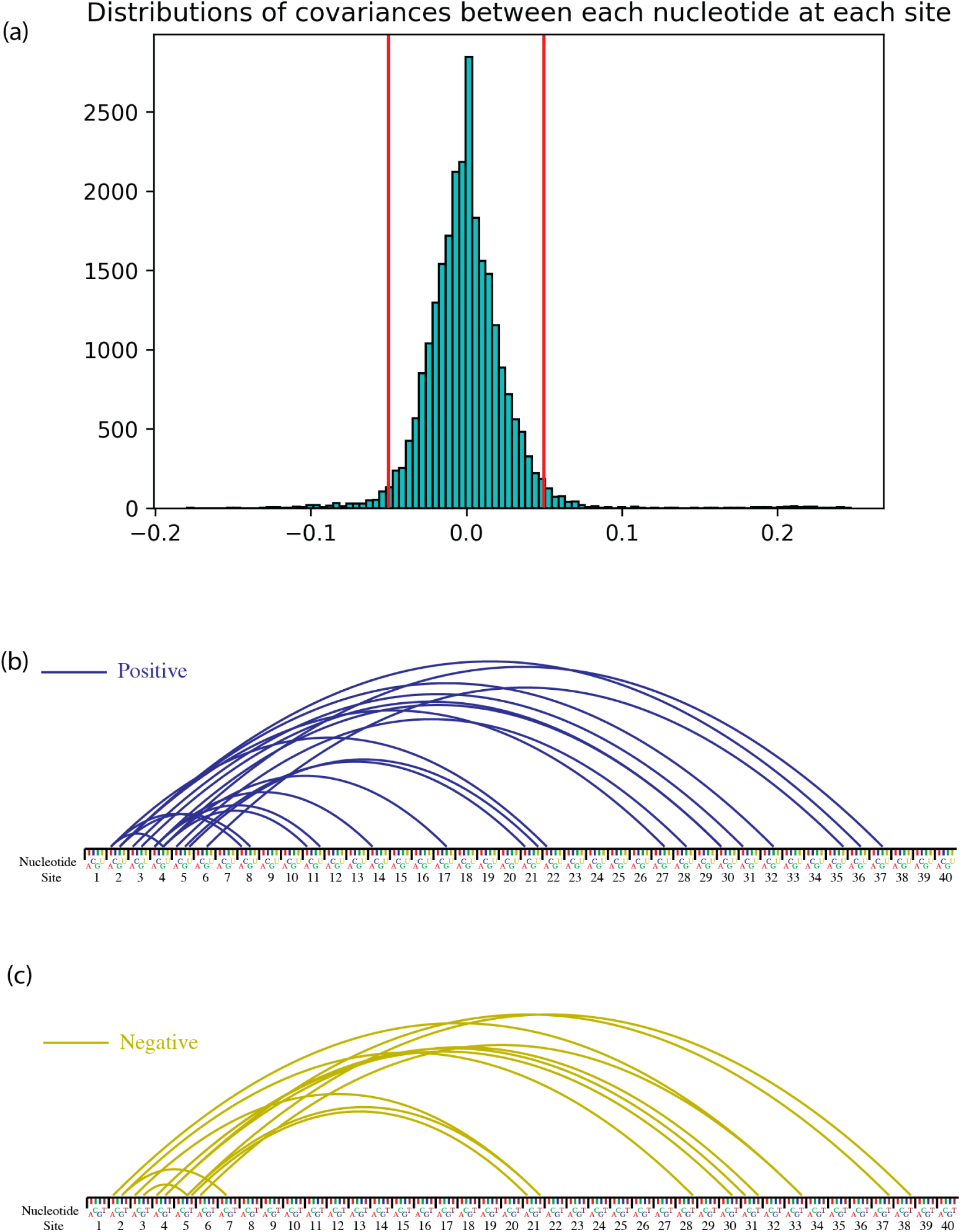
Strong correlations between nucleotides at sites across STAR. (a). Distribution of covariances between nucleotides at each site. Red bars indicate cutoffs of −0.05 and 0.05. (b) Large positive covariances with absolute magnitude greater than 0.05 across the 40 site long linear region of STAR (shown here from 5’ to 3’). (c) Large negative covariances. See Supplementary Table 1 for actual values.

Another possible explanation for this result could be related to the way these sequences were designed by the NUPACK algorithm. The objective of this software program is to calculate equilibrium distributions of the given nucleic acid strands (21). However, as has been noted by the authors of the STAR paper, STAR “regulation is governed by kinetic, out-of-equilibrium folding regimes” and thus, the NUPACK design of STAR sequences may not resemble natural sequences (19). Therefore, the apparent importance of sites at the 5’ end of the sequence could be an artifact of the software used to design the sequences. This warrants further investigation.

The covariation pattern here indicates the potential importance of sites 2-6 to the functionality of this sequence, however, the fact that these sites appear to be correlated with a lot of other sites might suggest that they’re interacting with those other sites. Therefore, a functional variable must be projected onto this sequence space to find out the relative importance of these sites. First, we conduct first order analysis and project the phenotype onto just individual sites to see which sites by themselves seem to be most important and what nucleotides at those sites seem to stand out. Subsequently, when doing second order analysis, we look at pairwise interactions to see how interactions between sites affect the phenotype.

### Regressions of fold and off values onto linear binding regions

In the STAR system, the authors of the STAR paper determined that the OFF state provides the best measure of the efficiency of the STAR:target complex as a whole. This motivates the analysis of the relationship between sequence and function of the target RNA (as measured by termination efficiency). To establish this, after building the first order orthogonal polynomials, we computed regressions of OFF values onto each nucleotide at each site of the target RNA. First order analysis shows the direct effects of nucleotides at each site on the phenotype. The regressions of the phenotype onto the first order orthogonal polynomial show how each nucleotide at each site contributes to the overall function of the STAR sequence. Figure 6 shows these regressions and captures the connection between sequence and expression when looking at just first order patterns.

#### Distribution of regressions

The magnitude of regressions is highly variable but large values seem to be uniformly distributed across the sequence. Regressions here refer to the projection of the phenotype (OFF fluorescence values) onto the first order conditional polynomial (see Methods). This indicates that this pattern is “unstructured” as noted by the authors of the STAR paper. This can be inferred from the shape of the regressions across the linear region of the target RNA (Figure 6(a)). Given the covariance structure in the STAR linear sequence sequence (which is complementary to the target linear region) (Figure 5), we expected the regressions onto the sites to show a similar pattern of connection between sites at the 3’ end of the sequence and sites at the 5’ end. However, this pattern of correlations does not emerge when projecting the OFF fluorescence values onto the sequence space. Though there exists a dynamic range of GFP expression in the STAR library, as indicated by the OFF fluorescence values, the sequence correlation structure identified earlier does not seem to be functionally relevant when this phenotype is projected onto the sequence space. This is evidenced by the fact that when looking at just each site on its own, no particular sets of sites stick out as being more important than other parts of the sequence. To investigate this further, we built second order orthogonal polynomials and projected the phenotype onto them in order to identify pairs of interacting sites and any potential long-range interactions.

#### Overrepresentation of purines

While there is no spatial pattern in the magnitudes of regression coefficients, as was originally expected due to the sequence covariance pattern, there is a strong relationship between the kind of nucleotide at a site and the regression of the off phenotype values on it. Almost all the purines have positive regressions while almost all the pyrimidines have negative values. This means that there is a preference for having purines along the target RNA sequence while pyrimidines are disfavored. The hairpin of the target RNA, the intrinsic terminator, includes a string of guanines and cytosines which implies that these nucleotides would not be preferred along the linear region (19). But then this would mean that As and Us would have positive regressions here while Gs and Cs would be disfavored and have negative values. But that’s not what we see, we see that Gs and As have positive regressions and Cs and Us have negative regressions. So this is a somewhat mysterious result that warrants further investigation.

After noting the purine/pyrimidine distinction, we were led to an additional hypothesis regarding molecular weights. There is a positive relationship between the magnitude of regression and the molecular weight of the nucleotide. Guanine is the heaviest molecule, followed by adenine, uracil and cytosine (Figure 6(b)). The regressions were the largest for heaviest molecules and decreased as the molecular weights decreased.

Furthermore, adenine and guanine are capable of hydrogen bonding with uracil. Guanine has a triple bond with cytosine and can form a double (wobbly) bond with uracil. As both of these nucleotides are capable of hydrogen bonding with uracil and this is a unique feature that distinguishes them from the pyrimidines, it might be the case that these nucleotides are more preferred.

#### Regressions of ON values onto STAR sites

In addition to projecting OFF values onto target RNA sites, ON values were projected onto the space of the sequences comprising the STAR linear region. The ON values are a measure of how good a STAR is at binding to the target RNA and activating the downstream gene. Since the STAR sequence is complementary to the target RNA, as expected, the regressions of having pyrimidines in the STAR linear region are positive while the regressions of having purines is negative (Figure 7).

### 2nd order analysis on all pairs of sites across the STAR sequence

The first order analysis, which includes calculating covariances and variances of nucleotides at each site, revealed a number of highly correlated sites (absolute values of covariances being greater than .05) sequestered in the 5’ region of STAR. In order to test the hypothesis that sites that are correlated (Figure 5) are interacting in their effect on the phenotype, we built second order polynomials for each pair of sites across the STAR sequence (with 780 unique pairs in total). See the corresponding supplementary section that shows an example of how this is done with a different set of sites.

After building second order polynomials for each pair of sites across the STAR sequence, regressions of the phenotype onto each pair of sites were computed. For a given pair of sites, this results in a 4×4 matrix, with 16 total values that each correspond to a given nucleotide at the first site and another nucleotide at the second site. Figure 8 shows an example of this for sites 3, 5 and 21. The regressions shown in this figure are scaled by the absolute value of the largest regression across all combinations of pairs in the sequence.

**Figure 6.**
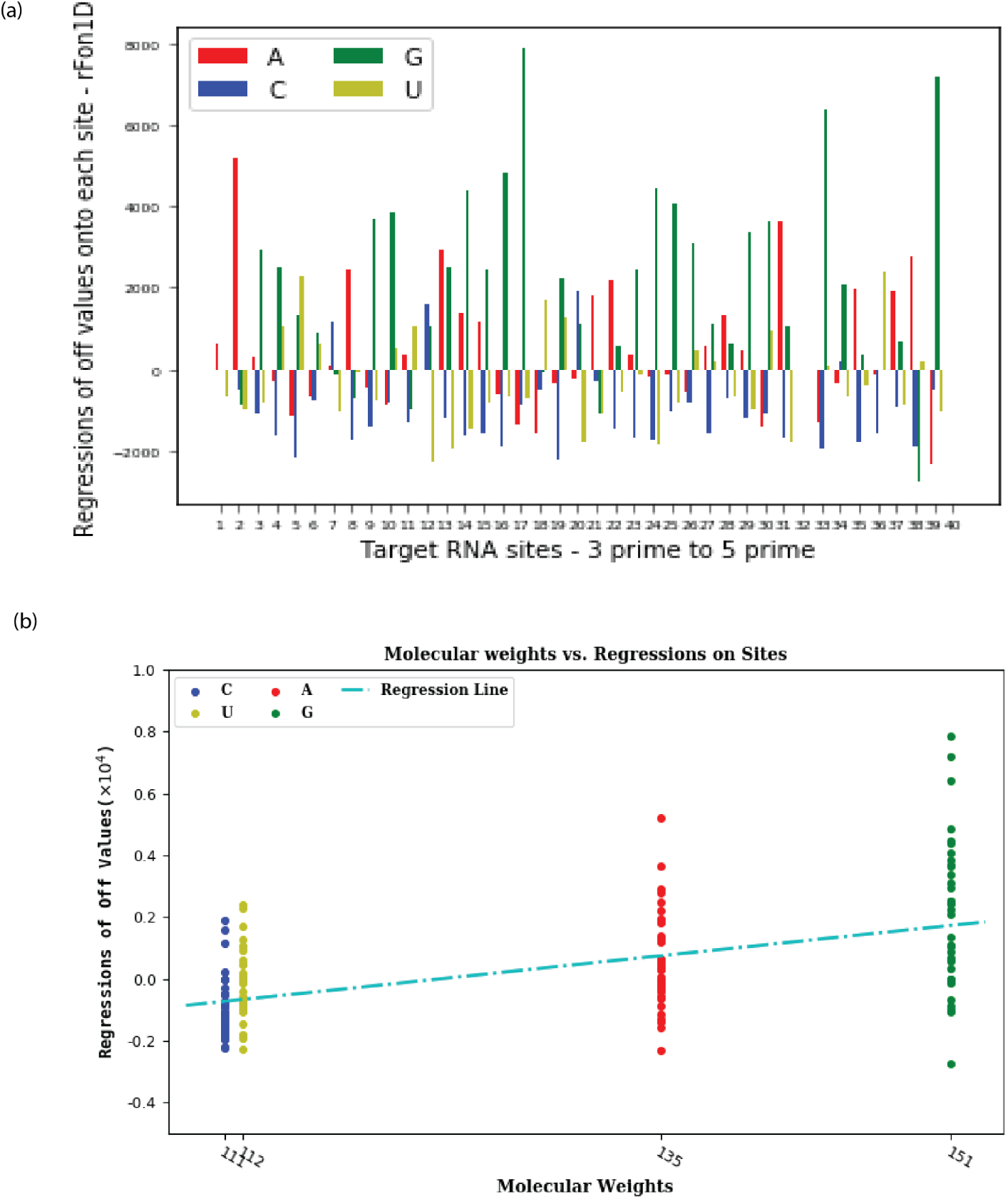
Regressions onto target RNA sites. (a) Regressions of off values onto each site of the target RNA orthogonalized within each vector, 3’ to 5’ 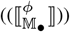. (b) Regressions of off values onto target RNA sites increase with increasing molecular weights of the nucleotides.

**Figure 7.**
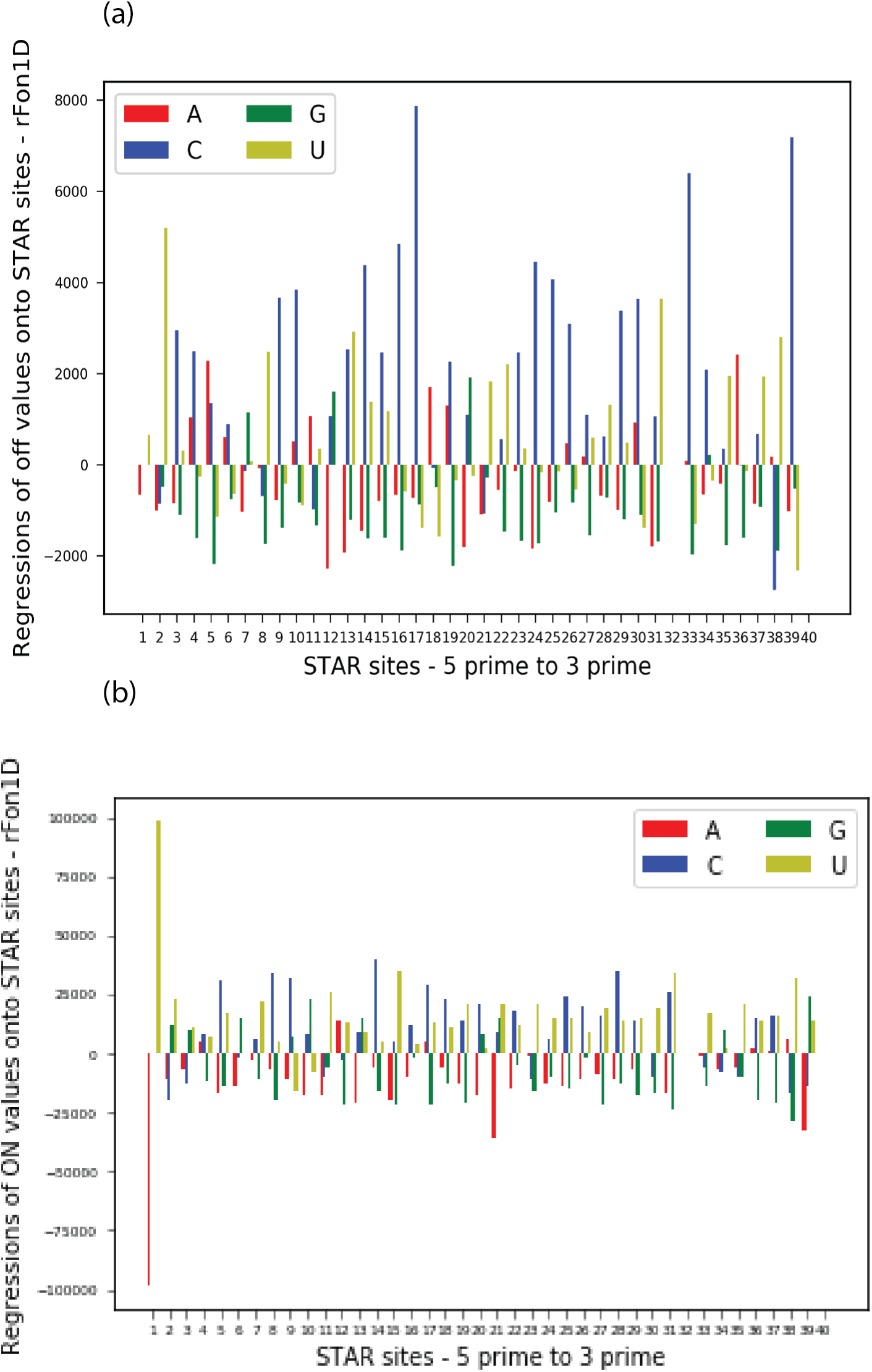
Regressions onto STAR sites. (a) Regressions of off values onto each STAR site orthogonalized within each vector (5’ to 3’). (b) Regressions of ON values onto STAR sites (orthogonalized within each vector).

**Figure 8.**
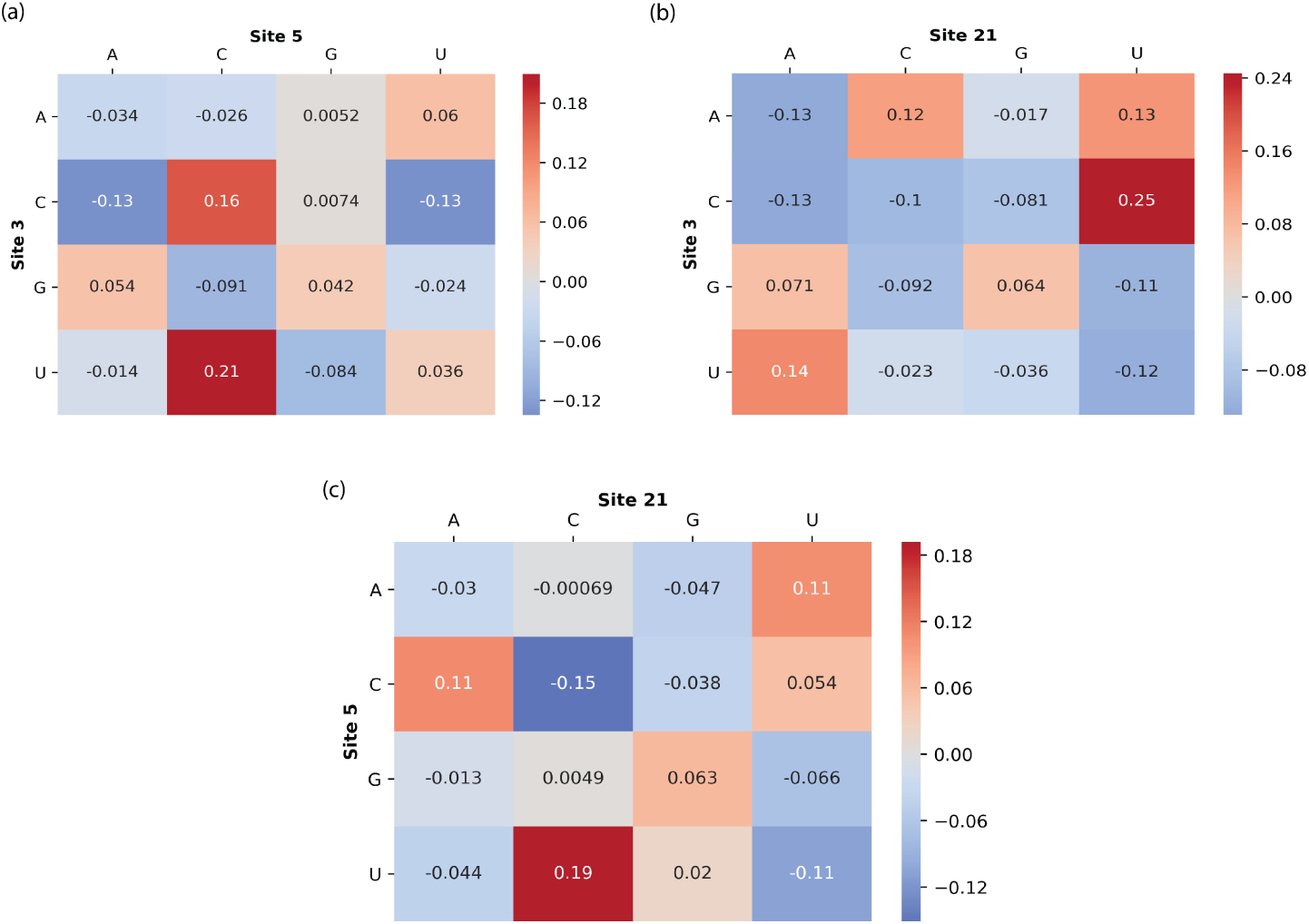
Regressions of off values onto the second order orthogonal polynomial when including the entire sequence. (a) shows regressions of OFF values on pairs of nucleotides, one at site 3 and one at site 5, independent of the first order contributions of each site. (b) shows regressions of OFF values on pairs of nucleotides at site 3 and at site 21. (c) shows regressions of OFF values on pairs of nucleotides at site 5 and at site 21. All values are scaled by the absolute value of the largest regression in the set of all combinations of pairs (i.e., 780 pairs formed by two sites across the 40 site long sequence).

To determine the effect of increasing distance between a pair of sites on the phenotype, we took the distances between the sites and plotted them against the absolute values of the maximum regressions (these are regressions of the phenotype onto two sites at a time). Figure 9(a) shows the sampling distribution of these regressions. It can be seen that a few regressions exist at the very tail end of the distribution as shown in the red part of the histogram. These regressions, all greater than 35,000, are also shown as a cloud of points, colored red, that appear in the top part of Figure 9(b). To visualize which sites made up these site pairs and which nucleotides at these sites correspond to the high positive or negative regressions, a plot similar to the covariance figure (Figure 5) was constructed (Figure 9(c)) with actual values given in Supplementary Table 2. For example, the blue curve connecting ‘A’ at site 5 with ‘C’ at site 15 means that having that combination contributes substantially to the OFF value, independently of the individual contributions of these sites by themselves. This visualization shows that there is no tendency for strongly interacting sites to be adjacent. In addition, there does not appear to be high regressions of the phenotype onto site pairs in the 5’ part of the sequence (as predicted by the covariance structure in Figure 5).

**Figure 9.**
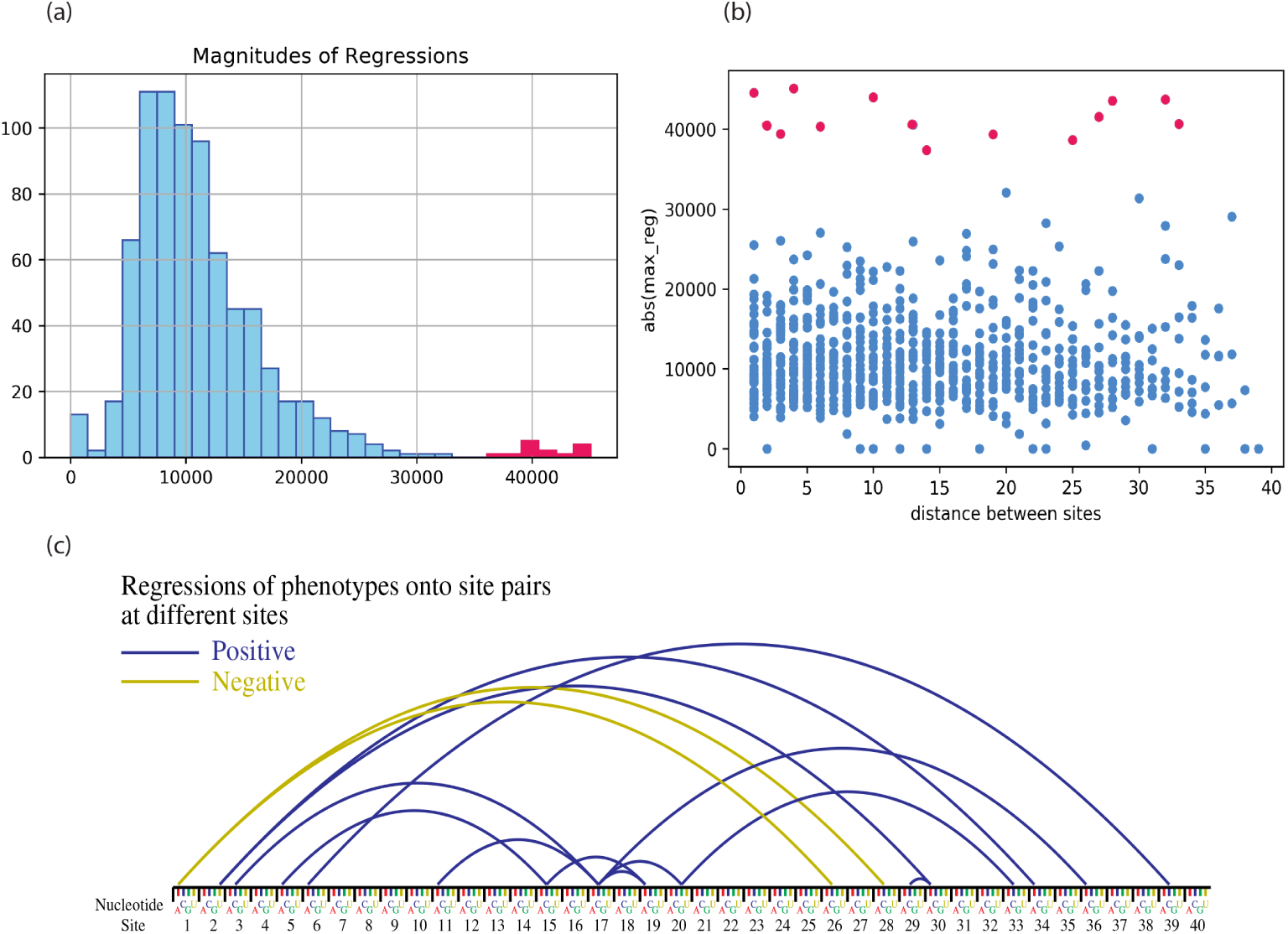
Regressions of the phenotype onto 780 site pairs along the STAR sequence. (a) shows the sampling distribution of absolute values of the largest regressions of the phenotype onto combination of two sites. Notable regressions are those at the tail end of the distribution, and are shown in red (> 35,000). (b) Relationship between distances across STAR sites and absolute values of the largest regressions with 14 site pairs with high regressions shown in red. (c) To further zoom into the 14 pairs of sites with notable regressions, as depicted in (a) and (b), curves are drawn between the interacting nucleotide at one site and the corresponding nucleotide at another site. This indicates a more global structure across the STAR sequence, instead of a case in which regressions of the phenotype onto pairs of sites in the 5’ region of the sequence are greater than those in the 3’ region.

Out of the 14 site pairs shown in Figure 9, there are 4 pairs that are between sites in the 5’ and 3’ regions. This potentially supports the hypothesis mentioned earlier which states that possible binding between these parts of the sequence would allow the hairpin region of the RNA to stay intact (by not binding with the hairpin) and pursue its function of transcription termination of the downstream gene. However, in addition to this long-range interaction, there are a cluster of strongly interacting sites in the middle portion of the sequence, between sites 11 and 20. This includes site 17 which has the most interactions (a ‘C’ at site 17 interacting with five other sites). There exists large values for regressions of the phenotype onto these site pairs (see Supplementary Table 2 for values). This suggests that there might exist some level of potential intramolecular binding in the middle section of the sequence that was not picked up by the NUPACK software when the initial sequences were being designed.

## DISCUSSION

In this work, we propose a novel mathematical tool to describe sites along biological sequences as vectors and quantify sequence-function relationships by projecting phenotypes onto the sequence space. Given a set of sequences and corresponding phenotypic data for each sequence, tensor-based orthogonal polynomials can be constructed based on the actual variation in sequences. The regression of phenotypes onto these polynomials can quantify not only the effects of different nucleotides at individual sites, but also the effect on phenotype of combinations of nucleotides at different sites. To show proof of concept, this method was applied to a case of synthetic RNA regulatory sequences, described in previously published work (19), that were experimentally constructed with the goal of identifying how sequence structure and design motifs affect RNA regulatory activity.

This method has some fundamental parallels with machine learning (ML) methods, such as neural networks, in that they share the very first step of one-hot encoding nucleotides and converting them to vectors/arrays. Figure 10 shows the similarities and differences between the two approaches. In the ML approach, sequence and corresponding phenotype data is fed into an algorithm (such as a deep neural net) after which the algorithm is trained to learn the relationship between the phenotype and the sequence (see (25) for example). In the mathematical approach, the sequence data is used to build a space within which we can conduct further mathematical analysis such as mapping phenotypes onto the sequence space. We argue that doing such mathematical analysis has some advantages because, here, the meaning of the values produced through this approach is clear because they were derived through the mathematics whereas when these results are derived from black box approaches, it is more difficult to interpret them. For example, in the analytical approach, we have a clear understanding of what a projection of a variable onto the polynomial space is, whereas this would be difficult to grasp in the traditional deep learning approaches.

**Figure 10.**
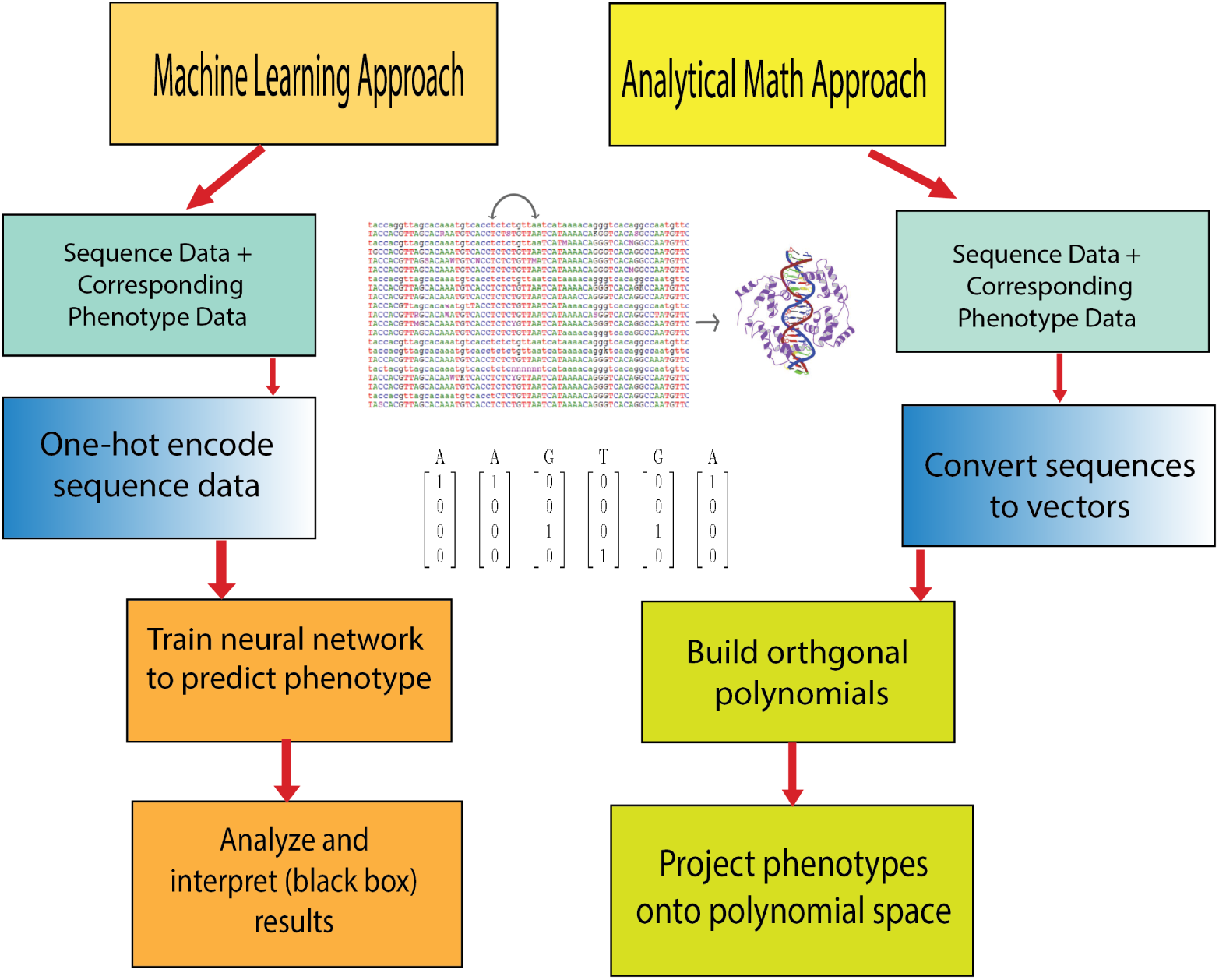
Comparison of analytical and ML approaches. Overview of similarities and differences between a machine learning approach and an analytical mathematics approach such as the method described here. The two share similar first steps and are ultimately aiming to understand sequence-phenotype relationships.

One other advantage that this approach has over other ML approaches is that once the sequence space is constructed (which is the part that requires most of the computational resources), one can easily project any corresponding phenotypes onto this space. In the ML approach, on the other hand, the algorithm would have to be re-trained entirely if one wanted to map a different phenotype to the underlying sequence. Furthermore, these two mappings would not be easily comparable as each phenotype and underlying sequence would have been trained differently. However, a hybrid approach that combines the two methods could be implemented. For example, it is possible to approximate some types of orthogonal polynomials using neural networks as has been done before (26, 27). A hybrid approach combining the tensor-based orthogonal polynomial method described here with a deep neural network trained on sequence and phenotype data is a promising area of further investigation.

Application of our method to the case of these regulatory RNAs showcase that even though a cluster of sites near the 5’ end are correlated with other sites throughout the sequence, there is no obvious preference for correlations concentrated at the ends of the sequence. It turned out that the correlation structure shown in Figure 5 did not have any functional relationship with the phenotype when the phenotype of OFF fluorescence values was regressed onto the sequence space. After identifying sites along the STAR sequence that are correlated with each other, we built second and third order orthogonal polynomials that quantified the effect of the phenotype on two-way and three-way combinations of sites (Supplementary Figures S5-S10).

In order to test whether the 5’ region contained combinations of sites that had a greater effect on the phenotype than those at the 3’ end, we built second order polynomials of combinations of two pairs of sites across the STAR sequence. Since the sequence is 40 sites long, there were 780 unique combinations. When projecting the phenotype onto these pairs of sites, we assessed the impact of distance between the pair of sites on the degree to which they influence phenotype, and whether combinations of sites in the 5’ region would have a greater effect on phenotype. As seen in Figure 9, instead of one region of the sequence being more important than the other, there seems to be a global structure to the interacting site pairs across the sequence that have high regressions.

The interaction data (Figure 9) does show some long range interactions, but it also shows a cluster of interacting sites in the middle. Furthermore, the pattern of interaction between sites in their impact on phenotype is not predicted by the nucleotide correlations between sites. Interactions at the 3’ and 5’ ends of the site, combined with those occurring in the middle of the sequence, suggest the possibility of competing intramolecular interactions and secondary structure formation in the absence of the target RNA. The authors utilized NUPACK aiming to minimize competing intramolecular interactions, however, our analyses indicate that there exist substantial competing intramolecular interactions that can interfere with the intermolecular interaction between the STAR:target complex. This aspect was not predicted sufficiently by the NUPACK algorithm. This presents an exciting opportunity to use novel computational and mathematical design approaches to inform experimental data and use the results to refine the design approach.

While direct Watson-Crick binding of the STAR to the target is a critical component of the STAR function, the function of this molecule will also likely be impacted by its potential to form secondary structures in the absence of the target RNA. Some level of secondary structure could be beneficial, perhaps by providing the STAR a level of stability from spontaneous degradation or nuclease-mediated degradation whereas other structures may negatively impact its stability for the same reasons. Additionally, a high amount of structural potential may create too much intramolecular binding and not allow for the intermolecular binding between the STAR and the target. However, for other reasons that might not be easily predicted, a certain level of STAR structure might be beneficial in the intermolecular binding.

## CONCLUSION

In conclusion, our methods provide a mathematical tool to find patterns in sequence data and to quantify the effect of the corresponding phenotype on the underlying sequence structure. Using this vector-based orthogonal polynomial approach, we can not only look at global patterns of sequence structure but can also identify the nucleotide state at each given site and how this affects phenotype at first and higher order levels.

Implementing this approach can be thought of as a two-step process. First, we analyze the existing covariance structure in the sequence data to see how different parts of the sequence might be correlated with one another (5). Second, we use the observed patterns of covariance to construct an orthogonal sequence space, into which we can project another variable to see which, if any, properties of the sequence predict the functional variable (Figures 7, 8, and 9).

In the case of the STAR sequence, the covariance analysis showed that a group of adjacent nucleotides near the 5’ end of the sequence were correlated with other nucleotides spread throughout the rest of the sequence. To see if this correlation corresponds to functional interactions between those nucleotides, we projected the experimental STAR fluorescence data into the orthogonal sequence space. This analysis identified a number of pairs of nucleotides that interact in their effect on fluorescence (Figure 9). However, these were not the pairs of nucleotides that were correlated with one another (Figure 5). The method thus allows us to tease apart structural properties of a sequence and functional interactions between different elements of that sequence. Given structural data, our approach can be utilized to yield insight into potential non-canonical base pairing interactions for different RNA families. And when combined with existing tools such as NUPACK, it can identify these interactions and thus allow for the construction of candidate sequences that can be used for downstream experimental analyses.

The limitations of this method largely lie with the computational power and memory required to compute polynomials for sequences with a large number of sites for which higher than second order analyses are sought. The proof of concept shown here consists of 99 sequences that are 40 sites long. However, a large number of sequences with a smaller number of sites might be more optimal for this method. A further area of investigation is potentially implementing a hybrid approach that utilizes a deep neural network to identify sites along the sequence that are functionally more important and then using just these sites as inputs to the tensor-based orthogonal polynomial approach described here. The existing program and the resulting command line tool is written in the Python programming language, however, efforts to optimize and parallelize the program using other languages such as C++ or Rust is one of the current objectives.

While we have given proof of concept of this approach using an example of regulatory RNAs, this method can be applied to other questions that aim to understand sequence-phenotype interactions such as transcription factor binding sites and how their sequence composition gives rise to different TF binding energies along with applications in synthetic biology that aim to understand the relationships between the underlying RNA sequence and corresponding secondary structure (25, 28).

## ACKNOWLEDGEMENTS

The authors would like to thank Saugato Rahman Dhruba, Olga Botvinnik, and Vijayanta Jain for their insightful comments and ideas. We’d like to thank the authors of the STAR RNA paper, including James Chappell, for doing the initial work that we used to apply our methods to and for stimulating discussions and comments. We’d like to acknowledge the TTU CISER program, the TTU Department of Biological Sciences, and Chan Zuckerberg Biohub for their support throughout the time this work was being done.

