## Supplementary Information for "Analyzing Genomic Data Using Tensor-Based Orthogonal Polynomials with Application to Synthetic RNAs"

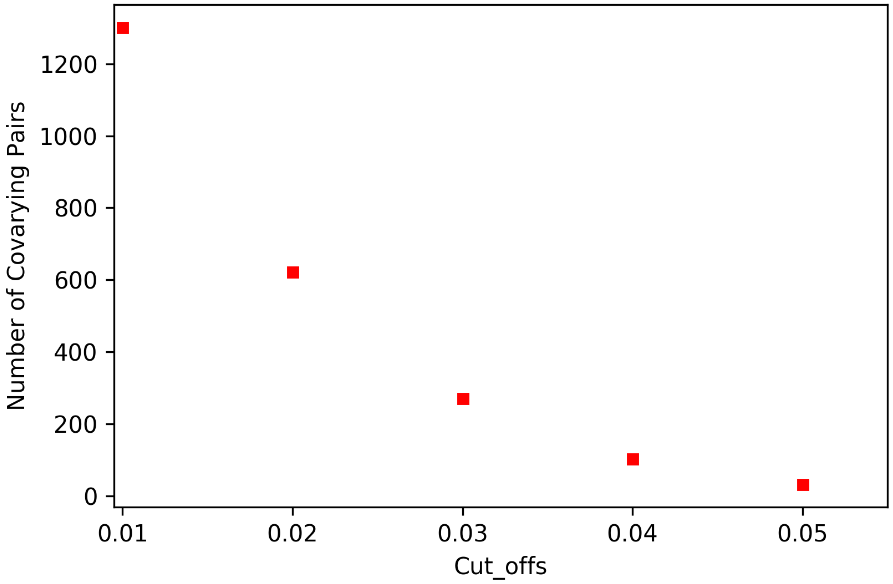


Figure 1: Determining a cutoff for highly covarying site pairs. This plot shows that the number of covarying site pairs drop off dramatically as the cutoff is increased.

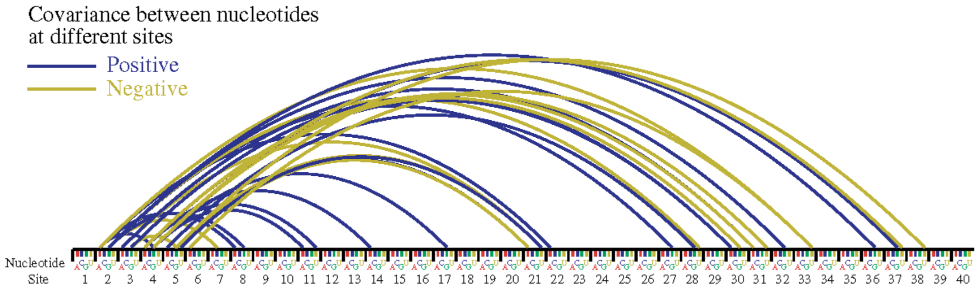


Figure 2: Set of 32 large positive and negative covariances (with cutoff of .05) between nucleotides at sites across the 40 site long linear region of STAR (shown here from 5' to 3').

Table 1: Covariances of nucleotides at given sites along the STAR sequence that are greater than 0.05 and less than $-0.05$.

| Site 1 | Site 2 | Nucleotide at Site 1 | Nucleotide at Site 2 | Covariance |
|---|---|---|---|---|
| 2 | 4 | A | G | 0.06325885 |
| 2 | 7 | G | C | -0.05142332 |
| 2 | 8 | A | A | 0.0580553 |
| 2 | 8 | G | G | 0.06152433 |
| 2 | 22 | A | A | 0.05387205 |
| 2 | 27 | G | G | 0.05325987 |
| 2 | 28 | G | U | -0.06336088 |
| 2 | 32 | G | G | 0.05693297 |
| 2 | 33 | A | U | -0.05305581 |
| 2 | 36 | G | G | 0.05325987 |
| 3 | 5 | G | G | -0.05428018 |
| 3 | 21 | A | U | -0.05662687 |
| 3 | 30 | G | A | 0.05978982 |
| 3 | 31 | A | A | 0.07284971 |
| 4 | 11 | A | U | 0.05540251 |
| 4 | 11 | G | A | 0.05081114 |
| 4 | 14 | A | A | 0.05315784 |
| 4 | 17 | G | G | 0.057035 |
| 4 | 30 | A | A | 0.05458627 |
| 4 | 30 | A | G | -0.05183145 |
| 4 | 31 | A | A | -0.05489236 |
| 4 | 31 | G | U | -0.05652484 |
| 4 | 37 | G | G | 0.06162636 |
| 4 | 37 | G | U | -0.05030099 |
| 5 | 21 | G | A | 0.05978982 |
| 5 | 21 | G | U | -0.05428018 |
| 5 | 21 | U | A | -0.0557086 |
| 5 | 21 | U | G | 0.05295378 |
| 5 | 28 | A | G | 0.05091317 |
| 5 | 38 | G | U | -0.05305581 |
| 6 | 33 | A | U | -0.05397408 |
| 6 | 35 | G | U | 0.05050505 |

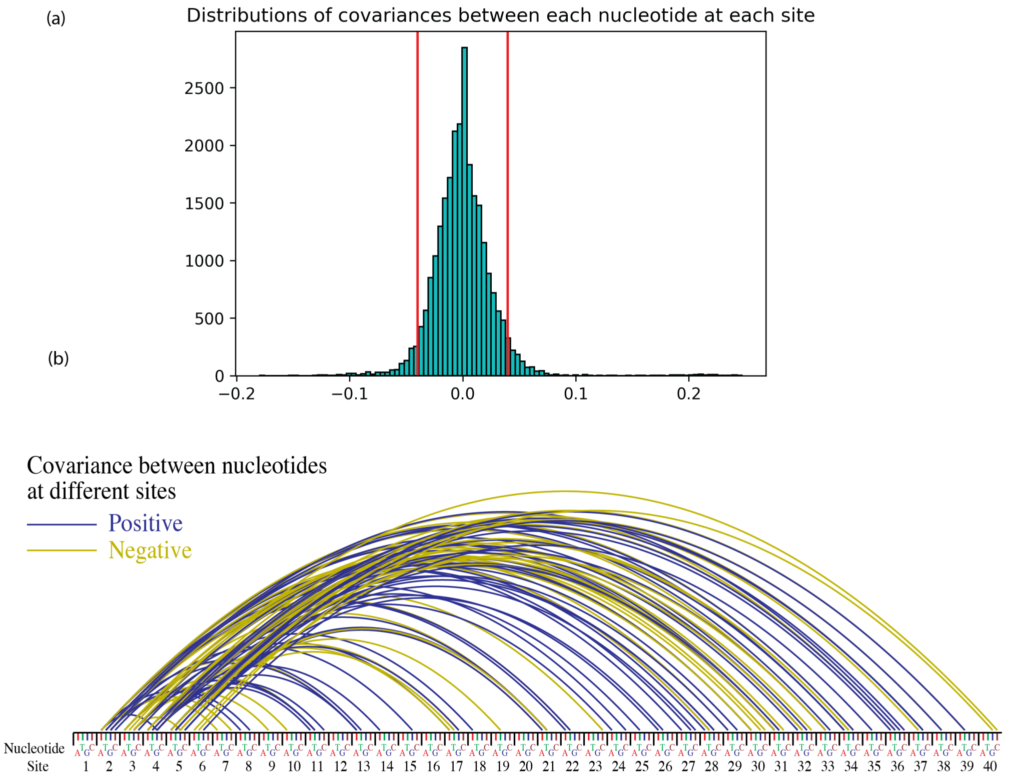


Figure 3: (a) Distribution of covariances between nucleotides at each site. Red bars indicate cutoffs of -0.04 and 0.04. (b) Set of 103 positive and negative covariances (with cutoff of .04) between nucleotides at sites across the 40 site long linear region of STAR (shown here from 5' to 3').

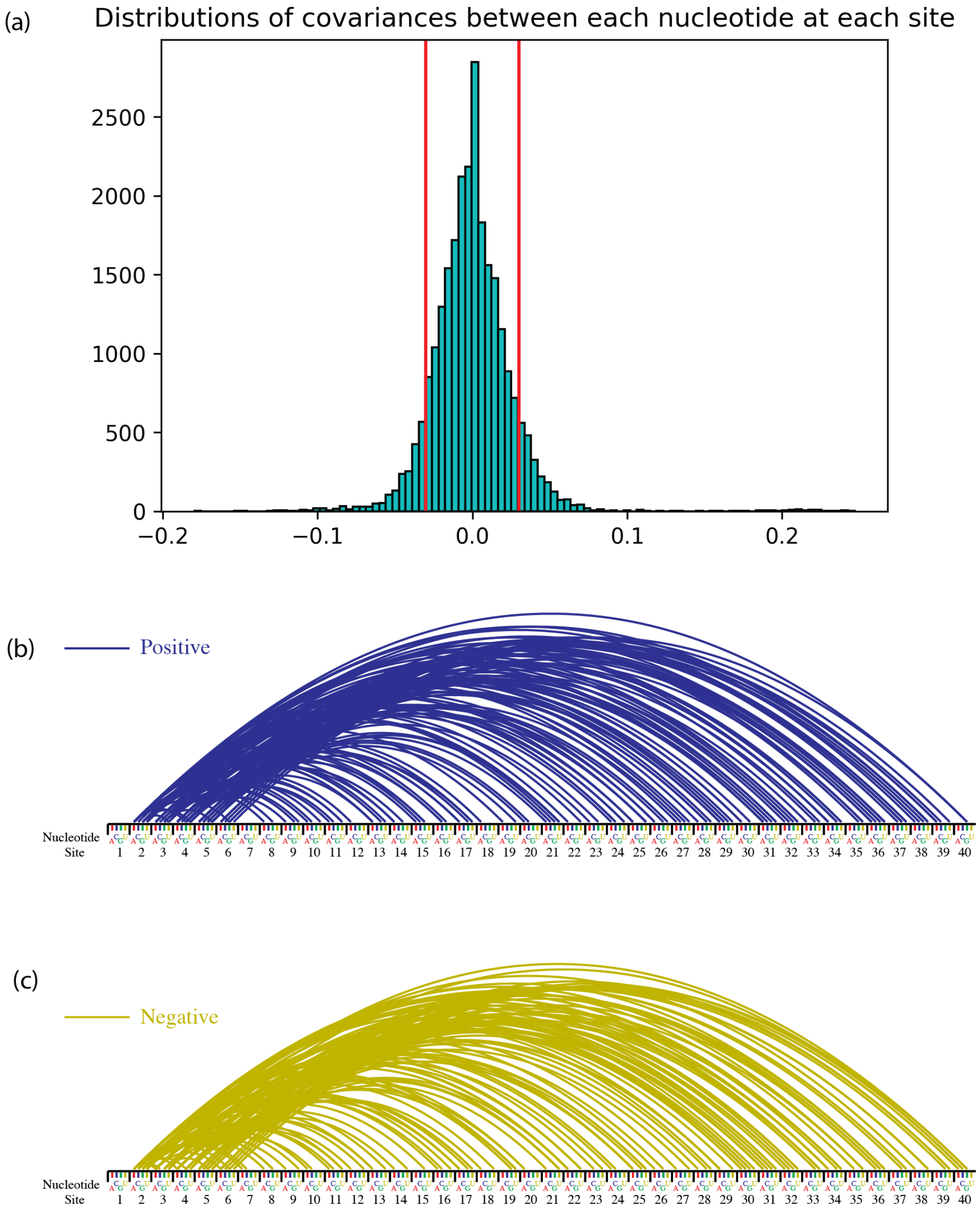


Figure 4: (a) Distribution of covariances between nucleotides at each site. Red bars indicate cutoffs of -0.03 and 0.03. (b) Set of 271 positive and negative covariances (with cutoff of .03) between nucleotides at sites across the 40 site long linear region of STAR (shown here from 5' to 3').

Table 2: Regression values corresponding to curves shown in Figure 9(c).

| Site 1 | Site 2 | Nucleotide at Site 1 | Nucleotide at Site 2 | Regression |
|---|---|---|---|---|
| 15 | 19 | C | A | 45080.62 |
| 29 | 30 | C | A | 44552.24 |
| 5 | 15 | A | C | 43990.82 |
| 2 | 34 | G | A | 43756.81 |
| 2 | 30 | G | A | 43568.87 |
| 1 | 28 | A | C | -41533.64 |
| 6 | 39 | A | C | 40675.03 |
| 20 | 33 | G | C | 40555.42 |
| 17 | 19 | C | A | 40456.85 |
| 11 | 17 | A | C | 40337.64 |
| 17 | 20 | C | G | 39410.58 |
| 17 | 36 | C | A | 39331.23 |
| 1 | 26 | A | C | -38653.73 |
| 3 | 17 | C | C | 37363.18 |

Table 3: Runtimes for building orthogonal polynomials for sequences and projecting phenotypes onto the sequence space. For the STAR/target example discussed in this work, first order analysis was done on all 40 sites. Second order analysis was done on two sites at a time and third order was done for a set of three sites (as discussed in the next section). The system used in this example was a Macbook Pro (2.9 GHz Intel Core i7) with 16 GB of memory. The program can run on Windows, Linux or Macintosh systems. See github repo for more information.

| Molecule | # of Sequences | Sites | Dimensions | Order | Time |
|---|---|---|---|---|---|
| DNA | 10,000 | 2 | 4 | 2nd | Approx. 35 mins |
| RNA | 99 | 40 | 4 | 1st | Approx 1.5 hrs |
| RNA | 99 | 2 | 4 | 2nd | 2 minutes |
| RNA | 99 | 3 | 4 | 3rd | 5 minutes |

### 0.1 Supplementary Section on 2nd and 3rd Order Examples

This section will go through examples of building second and third order polynomials given a set of interacting sites that have been determined *a priori*. This is to show the reader how such an analysis can be done. Note that this analysis is different from the analysis done for all 780 pairs of sites in the results section.

#### 0.1.1 2nd and 3rd order analyses on 3 covarying STAR sites

The first worked out example will go through a set of three sites for which we'll do second and third order analysis. As seen in Fig 5 and in Supplementary Table 1, it can be noted that site 3 covaried highly with site 5 and site 21 while site 5 also covaried with site 21. These covarying sites are where potential second order and third order interactions might be happening. To quantify this interaction, we can build the second and third order orthogonal polynomials for 3 sites (3,5,21) along the STAR linear region (from 5' to 3') and project off values onto this space (Figs S5 and S6).

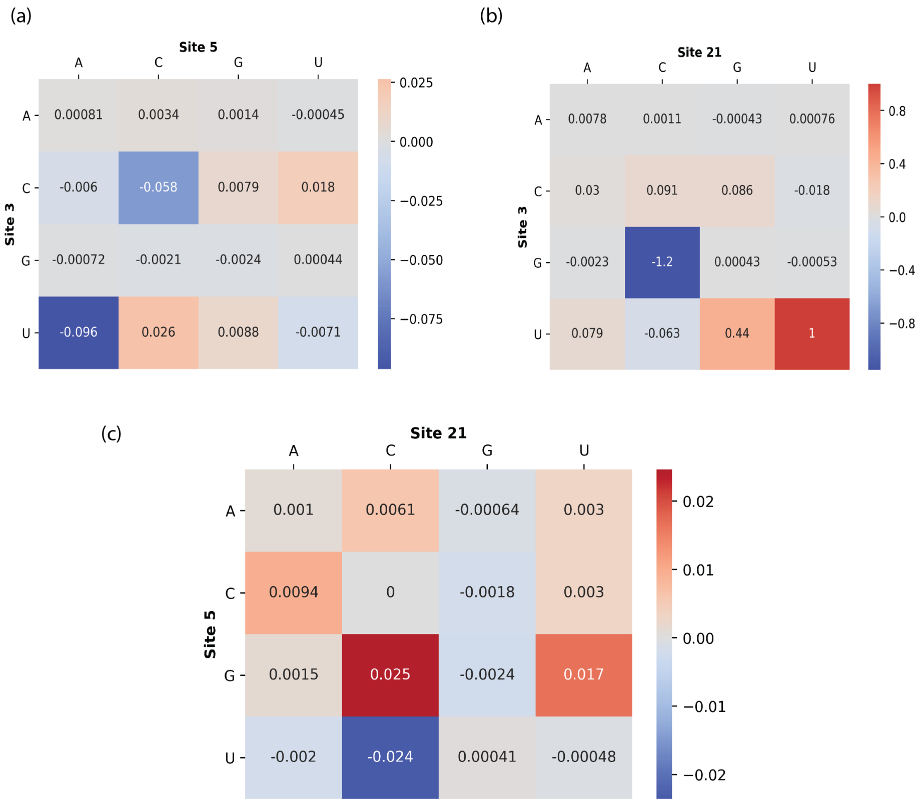


Figure 5: **Regressions of off values onto the second order orthogonal polynomial.** (a) shows regressions between having nucleotides at site 3 and those at site 5. (b) shows regressions between having nucleotides at site 3 and those at site 21. Here, the combination of a G at site 3 and a C at site 21 is highly disfavored while that of a U at site 3 and a U at site 21 is favored. (c) shows regressions between having nucleotides at site 5 and those at site 21. All values are scaled by the absolute value of the largest regression in the set of all combinations of pairs (i.e., pairs formed by sites 3 and 5, sites 3 and 21, and sites 5 and 21).

#### 0.1.2 Second order analysis on 6 covarying STAR sites

This second worked out example will go through a set of six sites for which we'll do second order analysis. To further assess the interacting sites sequestered in the 5' region of the STAR sequence and their effect on the phenotype, second order orthogonal polynomials can be constructed for the six sites (2,3,4,5,7, and 8). These six sites were chosen because they all are in the 5' region of the sequence and covaried highly with each other (Figure 5 and Supplementary Table 1). Since there are six sites, there are 15 unique pairs of interest (site 2 with sites 3,4,5,7,8, site 3 with sites 4,5,7,8, etc.). The regressions of off values onto the combinations of these sites are shown in Figs S7-S10. All values, across all 15 pairs of sites, are scaled by the absolute value of the largest regression in the set of these 15 two-way

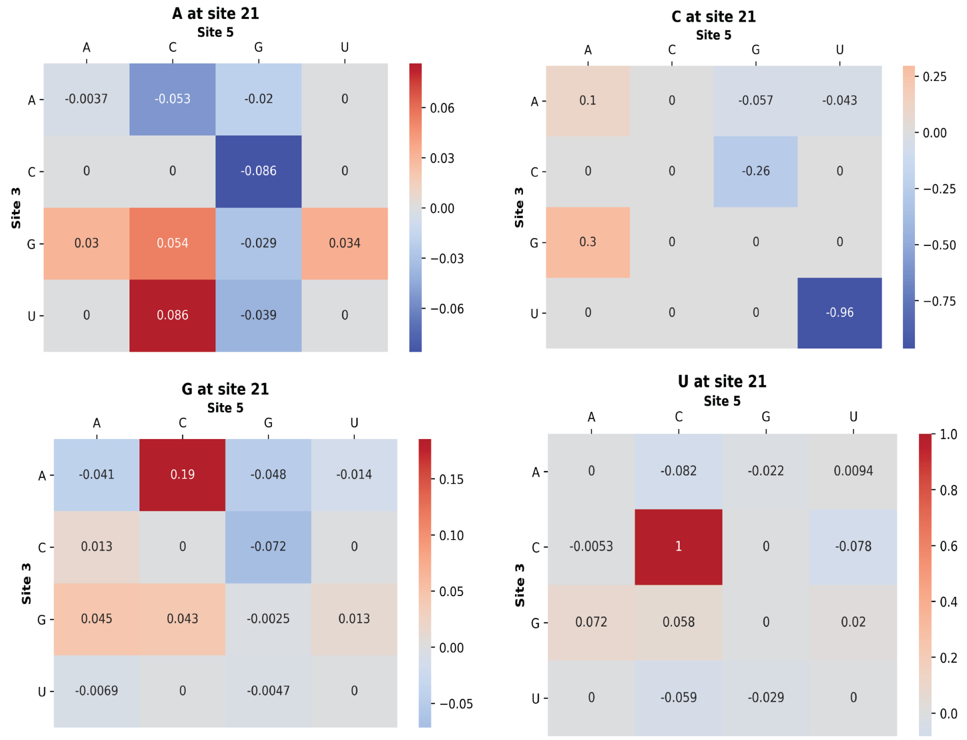


Figure 6: **Regressions of off values onto the third order orthogonal polynomial for three interacting sites.** All values are scaled by the absolute value of the largest regression in the set of all 3-way combinations of sites (64 total possibilities taking into account the different nucleotides at different sites).

combinations.

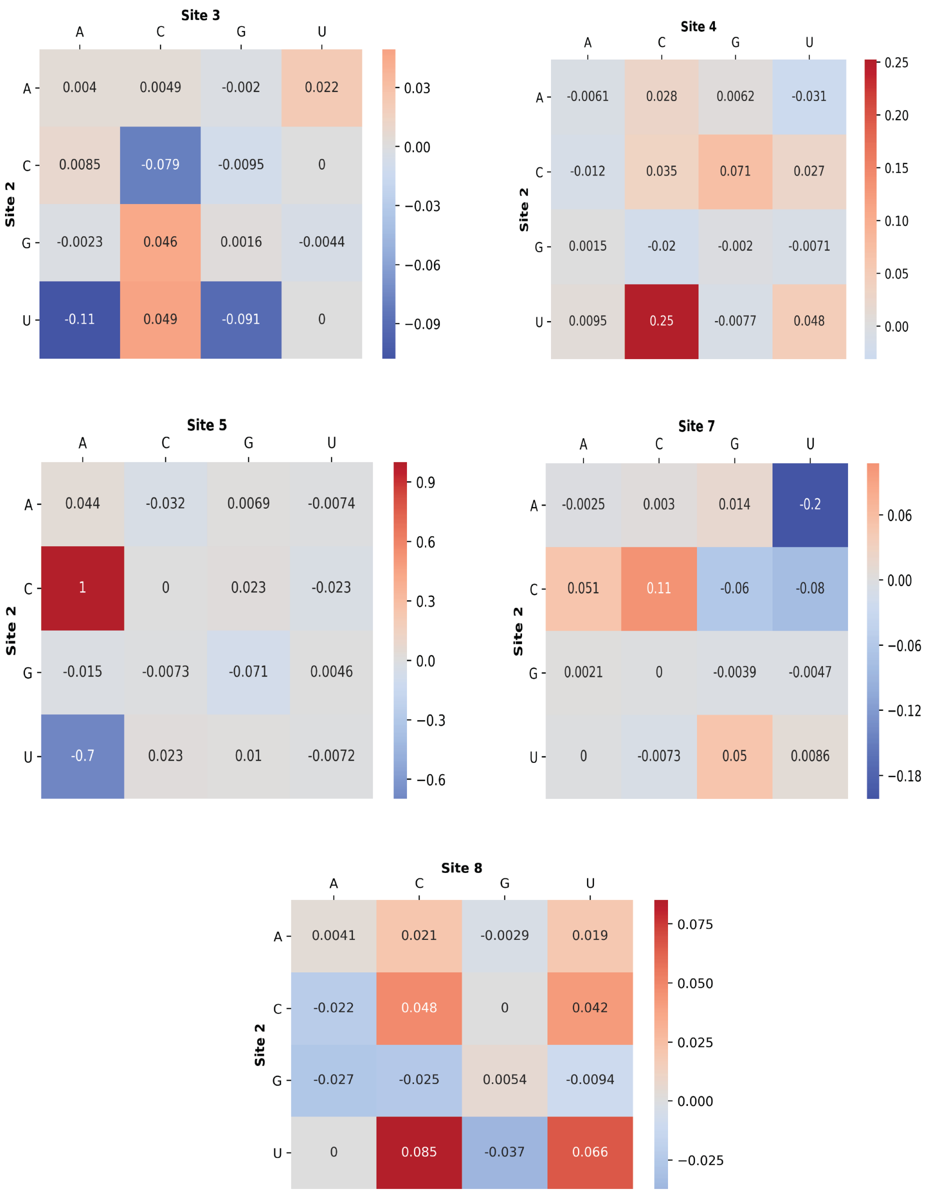


Figure 7: Regressions of OFF values onto combinations of site 2 with the rest of the 6 interacting sites.

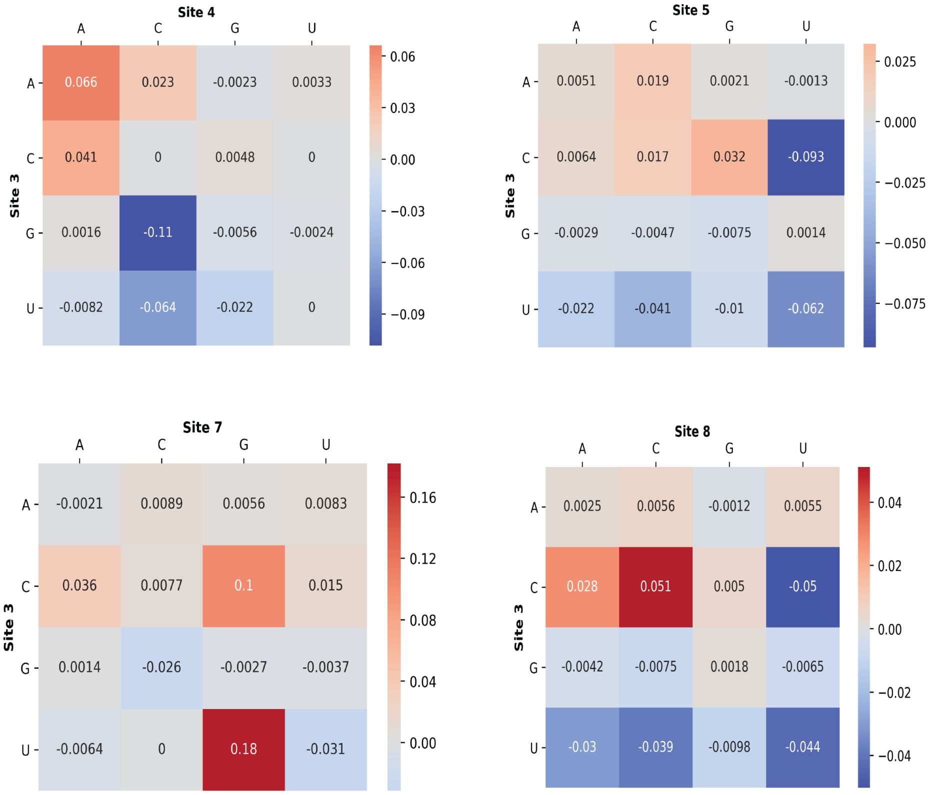


Figure 8: Regressions of OFF values onto combinations of site 3 with sites 4, 5, 7 and 8.

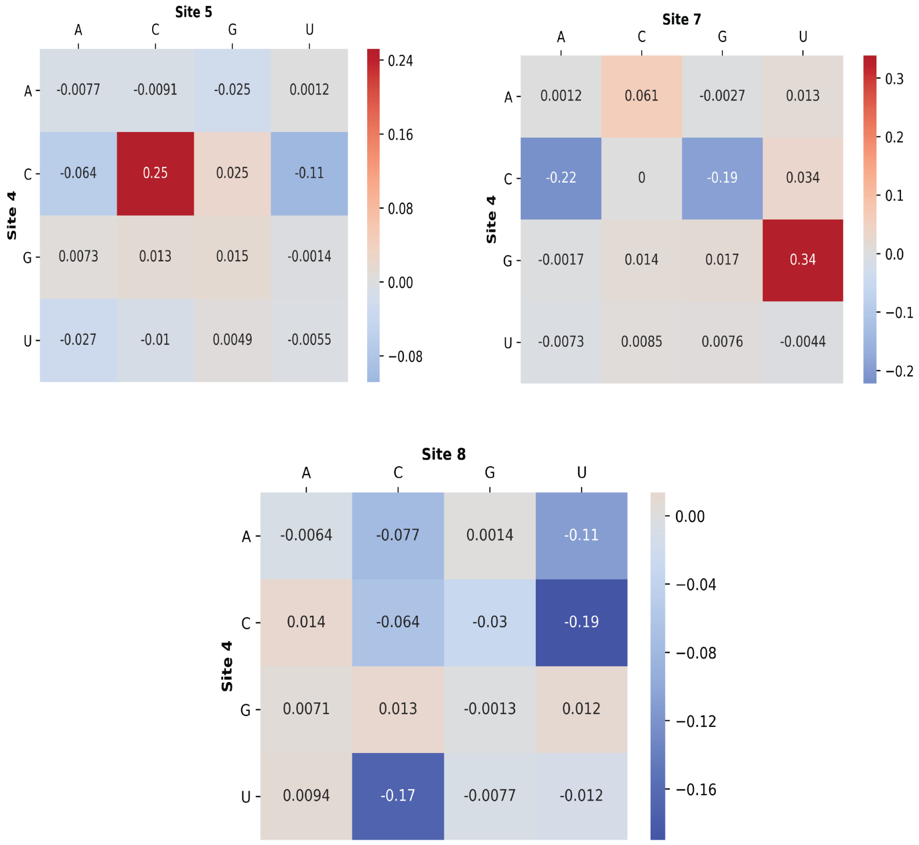


Figure 9: Regressions of OFF values onto combinations of site 4 with sites 5, 7 and 8.

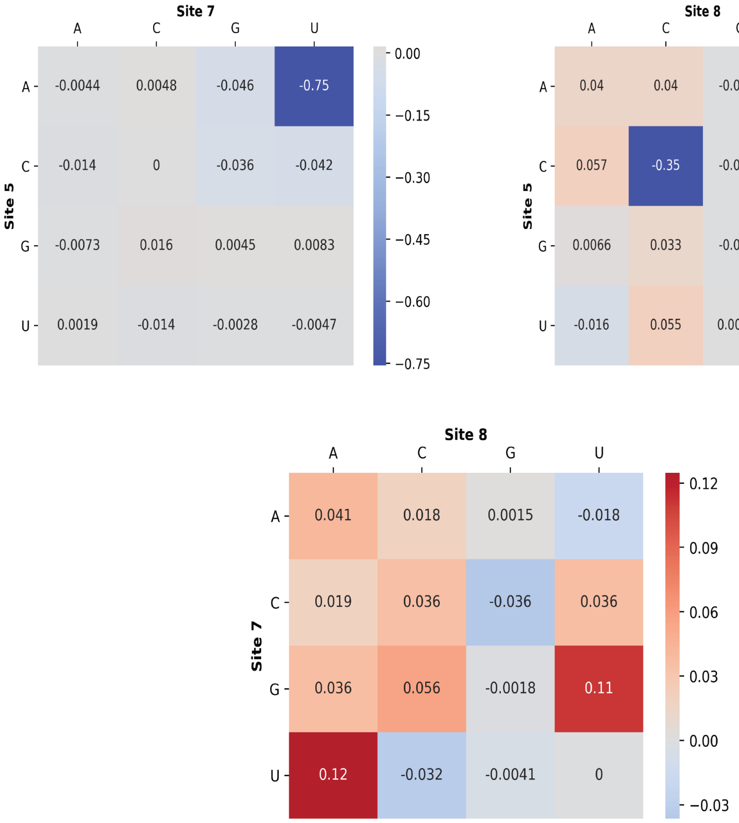


Figure 10: Regressions of OFF values onto combinations of sites 5 and 7, sites 5 and 8, and sites 7 and 8.
