## Supplementary Methods for "Analyzing Genomic Data Using Tensor-Based Orthogonal Polynomials with Application to Synthetic RNAs"

### Supplementary Methods - Orthogonal polynomials for phenotypes that are sequences

#### Scalar traits

For scalar traits (body mass, degree of altruism, etc.), we can write some other variable (fitness, susceptibility to a disease, *etc.*) as a function of the trait by constructing a set of orthogonal polynomials, of increasing order, for the trait. Orthogonality, here, is defined with respect to the distribution of variation in the population (*i.e.* the population distribution is the weight function). This is essentially generalized Fourier analysis.

For example, to write some variable $F$ as a function of a single phenotypic trait ($\phi$), we write the series:

$$F = \overline{F} + \sum_{i=1}^{N-1} [\![ {}^{F}_{\mathbb{P}^{[i]}} ]\!] \mathbb{P}^{[i]} \tag{1}$$

where $\mathbb{P}^{[i]}$ is the $i^{th}$ order orthogonal polynomial in $\phi$ (*i.e.* it has leading term $\phi^i$), $[\![ {}^{F}_{\mathbb{P}^{[i]}} ]\!]$ is the projection (regression) of $F$ onto polynomial $\mathbb{P}^{[i]}$, and $N$ is population size.

#### Sequences

Elements in a sequence (nucleotides, amino acids, *etc.*) are not accurately described by single scalar values (*i.e.* by a single number) because they are distinct types of things that are not distinuished by a single value. We thus represent monomers not with individual numbers, but with vectors.

For example: For a DNA sequence each nucleotide is represented by a 4 dimensional vector:

$$\begin{matrix} \text{A} & \text{C} & \text{G} & \text{T} \\ \begin{bmatrix} 1 \\ 0 \\ 0 \\ 0 \end{bmatrix} & \begin{bmatrix} 0 \\ 1 \\ 0 \\ 0 \end{bmatrix} & \begin{bmatrix} 0 \\ 0 \\ 1 \\ 0 \end{bmatrix} & \begin{bmatrix} 0 \\ 0 \\ 0 \\ 1 \end{bmatrix} \end{matrix}$$

One consequence of this is that the mean of a set of nucleotides is a vector that gives the distribution of nucleotides in the sample. For example, the mean of an A and a G would be:

$$\overline{(\text{A}, \text{G})} = \begin{bmatrix} \frac{1}{2} \\ 0 \\ \frac{1}{2} \\ 0 \end{bmatrix} \tag{2}$$

Note that this is essentially the same as the superposition of state vectors in quantum mechanics.

#### First order terms

To illustrate the approach, consider an example of four individuals, each with a two base sequence (the two bases that we are considering need not be adjacent to one another):

| Individual | 1 | 2 | 3 | 4 |
|---|---|---|---|---|
| Sequence | AC | CG | CC | GA |

We designate the $i^{th}$ monomer as $\mu_i$. The first order phenotype vectors for the four individual sequences above look like this:

Individual: 1 2 3 4

$\mu_1$ $\mu_2$ $\mu_1$ $\mu_2$ $\mu_1$ $\mu_2$ $\mu_1$ $\mu_2$

$$\left(\begin{bmatrix}1\\0\\0\\0\end{bmatrix}\begin{bmatrix}0\\1\\0\\0\end{bmatrix}\right)\left(\begin{bmatrix}0\\1\\0\\0\end{bmatrix}\begin{bmatrix}0\\0\\1\\0\end{bmatrix}\right)\left(\begin{bmatrix}0\\1\\0\\0\end{bmatrix}\begin{bmatrix}0\\1\\0\\0\end{bmatrix}\right)\left(\begin{bmatrix}0\\0\\1\\0\end{bmatrix}\begin{bmatrix}1\\0\\0\\0\end{bmatrix}\right) \tag{3}$$

As with polynomials of scalars, it is useful to subtract out the mean from each value (unlike the scalar case, though, we will continue to have use for the vectors in 3). Subtracting the mean yields the first order $\mathbb{M}$ vectors:

$\mathbb{M}^1$ $\mathbb{M}^2$ $\mathbb{M}^1$ $\mathbb{M}^2$ $\mathbb{M}^1$ $\mathbb{M}^2$ $\mathbb{M}^1$ $\mathbb{M}^2$

$$\left(\begin{bmatrix}\frac{3}{4}\\-\frac{1}{2}\\-\frac{1}{4}\\0\end{bmatrix}\begin{bmatrix}-\frac{1}{4}\\\frac{1}{2}\\-\frac{1}{4}\\0\end{bmatrix}\right)\left(\begin{bmatrix}-\frac{1}{4}\\\frac{1}{2}\\-\frac{1}{4}\\0\end{bmatrix}\begin{bmatrix}-\frac{1}{4}\\-\frac{1}{2}\\\frac{3}{4}\\0\end{bmatrix}\right)\left(\begin{bmatrix}-\frac{1}{4}\\\frac{1}{2}\\-\frac{1}{4}\\0\end{bmatrix}\begin{bmatrix}-\frac{1}{4}\\\frac{1}{2}\\-\frac{1}{4}\\0\end{bmatrix}\right)\left(\begin{bmatrix}-\frac{1}{4}\\-\frac{1}{2}\\\frac{3}{4}\\0\end{bmatrix}\begin{bmatrix}\frac{3}{4}\\-\frac{1}{2}\\-\frac{1}{4}\\0\end{bmatrix}\right)$$

The matrix of covariances for the two sites is just the mean, across all individuals, of the outer product of $\mathbb{M}^1$ and $\mathbb{M}^2$:

$$[\![\mu_1, \mu_2]\!] = \overline{\mathbb{M}^1 \otimes \mathbb{M}^2} = \begin{bmatrix} -\frac{1}{16} & \frac{1}{8} & -\frac{1}{16} & 0 \\ -\frac{1}{8} & 0 & \frac{1}{8} & 0 \\ \frac{3}{16} & -\frac{1}{8} & -\frac{1}{16} & 0 \\ 0 & 0 & 0 & 0 \end{bmatrix} \tag{4}$$

We can similarly find the variances of the elements of the $\mathbb{M}^1$ vectors. In this example they are:

$$
\begin{aligned}
[\![{}^2\mathbb{M}^1(0)]\!] &= \frac{3}{16} \\
[\![{}^2\mathbb{M}^1(1)]\!] &= \frac{1}{4} \\
[\![{}^2\mathbb{M}^1(2)]\!] &= \frac{3}{16} \\
[\![{}^2\mathbb{M}^1(3)]\!] &= 0
\end{aligned} \tag{5}
$$

#### Projections onto $\mathbb{M}$ coordinates

We now need to construct orthogonal polynomials based on our vectors. This turns out to be similar, but not exactly the same, as using polynomials based on scalars rather than vectors.

As shown in Equation 1, the projection of a variable $F$ onto a scalar based polynomial $\mathbb{P}^i$ is just the regression of $F$ on $\mathbb{P}^i$ (denoted $[\![\frac{F}{\mathbb{P}^i}]\!]$) multiplied by $\mathbb{P}^i$. For a vector based polynomial, $\mathbb{M}^i$, the projection of $F$ for a particular individual is given by:

$$
[\![\tfrac{F}{\mathbb{M}^i}]\!] \cdot (\mu_i \otimes \mu_i) \cdot \mathbb{M}^i \tag{6}
$$

Where "$\cdot$" represents inner product and "$\otimes$" represents outer product. The role of the new term, $\mu_i \otimes \mu_i$, is developed in the section **"Deriving the vector valued equation from the scalar case"** below. The intuitive explanation is that the elements within a single $\mathbb{M}^1$ vector are not independent of one another. To see why, go back to the $\mu$ vectors. If there is a 1 in a particular spot in a $\mu$ vector, then the other spots must be 0's. Though it is not as visually obvious, the same negative correlation between elements holds for the $\mathbb{M}$ vectors. We thus need to orthogonalize within each vector. It turns out that doing this causes all of the elements other than the one that was a '1' in the original $\mu$ vector to become 0; and this is exactly what we get when we inner multiply by $(\mu_1 \otimes \mu_1)$. As an example, this is what we get for the $\mathbb{M}^1$ vector of the first individual listed above:

$$
(\mu_1 \otimes \mu_1) \cdot \mathbb{M}^1 = \begin{bmatrix} 1 & 0 & 0 & 0 \\ 0 & 0 & 0 & 0 \\ 0 & 0 & 0 & 0 \\ 0 & 0 & 0 & 0 \end{bmatrix} \cdot \begin{bmatrix} \frac{3}{4} \\ -\frac{1}{2} \\ -\frac{1}{4} \\ 0 \end{bmatrix} = \begin{bmatrix} \frac{3}{4} \\ 0 \\ 0 \\ 0 \end{bmatrix} \tag{7}
$$

In calculating the regressions, we have to use the entire vector (not multiplied by $(\mu_i \otimes \mu_i)$), since there we are interested in the impact that each possible nucleotide at that site would have on $F$. When calculating the contribution of a site in a particular individual, the $\mu_i \otimes \mu_i$ term essentially prevents us from adding in the effects of nucleotides that are not present at the site in that individual.

Since the $\mathbb{M}^1$ and $\mathbb{M}^2$ vectors are not orthogonal, we need to construct a polynomial in $\mathbb{M}^2$ that is orthogonal to $\mathbb{M}^1$. We will denote this as $\mathbb{M}^2_1$, or as $\mathbb{M}^2_\bullet$ if there are only two sites.

Using Gram-Schmidt orthogonalization, we fine the polynomial for site 2 independent of site 1 as:

$$\mathbb{M}^2_1 = \mathbb{M}^2 - \left[\!\!\left[\frac{\mathbb{M}^2}{\mathbb{M}^1}\right]\!\!\right] \cdot (\mu_1 \otimes \mu_1) \cdot \mathbb{M}^1 \tag{8}$$

In calculating $\left[\!\!\left[\frac{\mathbb{M}^2}{\mathbb{M}^1}\right]\!\!\right]$, we take the regression between each element of each vector separately. This yields a matrix that, for the example system shown above, looks like this:

$$\left[\!\!\left[\frac{\mathbb{M}^2}{\mathbb{M}^1}\right]\!\!\right] = \begin{bmatrix} -\frac{1}{3} & \frac{2}{3} & -\frac{1}{3} & 0 \\ -\frac{1}{2} & 0 & \frac{1}{2} & 0 \\ 1 & -\frac{2}{3} & -\frac{1}{3} & 0 \\ 0 & 0 & 0 & 0 \end{bmatrix} \tag{9}$$

The elements of the matrix in Equation 9 are found by dividing the elements of the matrix of covariances in Equation 4 by the appropriate variance terms from Equations 6. If the variance is zero, then the corresponding covariance will also be zero and we define the regression as zero. (Note that the possibility of zero variances means that we can not write this using typical linear algebra methods, since we would have to invert a variance matrix that will often be singular.)

Note that the second term on the righthand side of Equation 8 is the inner product of a matrix and a vector, which yields a vector that can be added to the first term. Using this approach, we get the following orthogonal vectors for site 1 and for site 2 independent of 1:

$$\left(\overset{\mathbb{M}^1}{\begin{bmatrix} \frac{3}{4} \\ -\frac{1}{2} \\ -\frac{1}{4} \\ 0 \end{bmatrix}} \overset{\mathbb{M}^2_\bullet}{\begin{bmatrix} 0 \\ 0 \\ 0 \\ 0 \end{bmatrix}}\right) \left(\overset{\mathbb{M}^1}{\begin{bmatrix} -\frac{1}{4} \\ \frac{1}{2} \\ -\frac{1}{4} \\ 0 \end{bmatrix}} \overset{\mathbb{M}^2_\bullet}{\begin{bmatrix} 0 \\ -\frac{1}{2} \\ \frac{1}{2} \\ 0 \end{bmatrix}}\right) \left(\overset{\mathbb{M}^1}{\begin{bmatrix} -\frac{1}{4} \\ \frac{1}{2} \\ -\frac{1}{4} \\ 0 \end{bmatrix}} \overset{\mathbb{M}^2_\bullet}{\begin{bmatrix} 0 \\ \frac{1}{2} \\ -\frac{1}{2} \\ 0 \end{bmatrix}}\right) \left(\overset{\mathbb{M}^1}{\begin{bmatrix} -\frac{1}{4} \\ -\frac{1}{2} \\ \frac{3}{4} \\ 0 \end{bmatrix}} \overset{\mathbb{M}^2_\bullet}{\begin{bmatrix} 0 \\ 0 \\ 0 \\ 0 \end{bmatrix}}\right) \tag{10}$$

We can now project a scalar value, such as a measure of protein folding efficiency, into this basis as:

$$F = \overline{F} + \left[\!\left[\frac{F}{\mathbb{M}^1}\right]\!\right] \cdot (\mu_1 \otimes \mu_1 \cdot \mathbb{M}^1) + \left[\!\left[\frac{F}{\mathbb{M}^2_\bullet}\right]\!\right] \cdot (\mu_2 \otimes \mu_2 \cdot \mathbb{M}^2_\bullet) \tag{11}$$

As we did in building the orthogonal basis, we need to take the regression of $F$ onto each element of $\mathbb{M}^1$ or $\mathbb{M}^2_\bullet$. Since $F$ is a scalar, this yields a regression vector:

$$\left[\!\left[\frac{F}{\mathbb{M}^1}\right]\!\right] = \begin{bmatrix} \left[\!\left[\frac{F}{\mathbb{M}^1(1)}\right]\!\right] \\ \left[\!\left[\frac{F}{\mathbb{M}^1(2)}\right]\!\right] \\ \left[\!\left[\frac{F}{\mathbb{M}^1(3)}\right]\!\right] \\ \left[\!\left[\frac{F}{\mathbb{M}^1(4)}\right]\!\right] \end{bmatrix} \tag{12}$$

These vectors are inner multiplied by the vectors $\mu_1 \otimes \mu_1 \cdot \mathbb{M}^1$ and $\mu_2 \otimes \mu_2 \cdot \mathbb{M}^2_\bullet$, which for our example look like this:

$$\left(\begin{bmatrix}\frac{3}{4}\\0\\0\\0\end{bmatrix}\begin{bmatrix}0\\0\\0\\0\end{bmatrix}\right)\left(\begin{bmatrix}0\\\frac{1}{2}\\0\\0\end{bmatrix}\begin{bmatrix}0\\0\\\frac{1}{2}\\0\end{bmatrix}\right)\left(\begin{bmatrix}0\\\frac{1}{2}\\0\\0\end{bmatrix}\begin{bmatrix}0\\\frac{1}{2}\\0\\0\end{bmatrix}\right)\left(\begin{bmatrix}0\\0\\\frac{3}{4}\\0\end{bmatrix}\begin{bmatrix}0\\0\\0\\0\end{bmatrix}\right) \tag{13}$$

Note again that multiplying by the outer product of the $\mu$'s just has the effect of making all elements zero except those in the spot originally occupied by the '1' in the $\mu$ vectors. The fact that we regress on the vectors in 10, but then multiply this regression by the vectors in 13, is the main difference between working with polynomials of vectors *vs.* polynomials of scalars.

### Deriving the vector valued equation from the scalar case

In this section, we discuss how to go from treating each possible monomer as a distinct scalar-valued trait to treating the site as a vector-valued trait. This will also clarify the origin of the $\mu \otimes \mu$ terms in the equations above.

Consider a simplified genome within which each site can have one of three nucleotides (or monomers), $A$, $B$, or $C$ (the extension to 4 or more possible monomers is straightforward). We begin by treating each nucleotide at each site as a distinct trait, with a value of 1 if that nucleotide is present at that site, and 0 if it is not. Since we are now dealing with scalar traits, we denote them by $\phi$ and use $\mathbb{P}$ for orthogonal polynomials in $\phi$. For two sites each with three possible nucleotides, this would give us 6 different traits:

| | | | |
|---|---|---|---|
| $\phi_1$ | $A$ at site 1 | $\phi_4$ | $A$ at site 2 |
| $\phi_2$ | $B$ at site 1 | $\phi_5$ | $B$ at site 2 |
| $\phi_3$ | $C$ at site 1 | $\phi_6$ | $C$ at site 2 |

If the actual sequence were $BC$ ($B$ at site 1 and $C$ at site 2), then we would have $\phi_2 = 1$ and $\phi_6 = 1$, with all other traits being zero. This seems like a very unnatural way to represent a sequence, since it ignores which monomers are at which site, but we will see that we can collapse terms in such a way as to bring the identity of sites back into focus.

Construct orthogonal polynomials in the $\phi$ terms:

$$\begin{aligned} \mathbb{P}^i &= \phi_i - \overline{\phi_i} \\ \mathbb{P}^j_i &= \text{Polynomial in } \phi_j \text{ orthogonal to } \mathbb{P}^i, \quad \text{with } \overline{\mathbb{P}^j_i} = 0 \end{aligned} \tag{14}$$

Since there are many ways to construct the orthogonal polynomials, there are many ways that we can project $F$ into our sequence space. Equation 15 shows three such projections for the case of first order terms.

$$
\begin{aligned}
F &= \overline{F} + [\![\tfrac{F}{\mathbb{P}^1}]\!]\,\mathbb{P}^1 + [\![\tfrac{F}{\mathbb{P}^2_1}]\!]\,\mathbb{P}^2_1 + [\![\tfrac{F}{\mathbb{P}^3_{12}}]\!]\,\mathbb{P}^3_{12} + [\![\tfrac{F}{\mathbb{P}^4_{[3]}}]\!]\,\mathbb{P}^4_{[3]} + [\![\tfrac{F}{\mathbb{P}^5_{[3]4}}]\!]\,\mathbb{P}^5_{[3]4} + [\![\tfrac{F}{\mathbb{P}^6_{[3]45}}]\!]\,\mathbb{P}^6_{[3]45} + {2^{nd} \atop \text{order}} \\
&= \overline{F} + [\![\tfrac{F}{\mathbb{P}^2}]\!]\,\mathbb{P}^2 + [\![\tfrac{F}{\mathbb{P}^1_2}]\!]\,\mathbb{P}^1_2 + [\![\tfrac{F}{\mathbb{P}^3_{12}}]\!]\,\mathbb{P}^3_{12} + [\![\tfrac{F}{\mathbb{P}^5_{[3]}}]\!]\,\mathbb{P}^5_{[3]} + [\![\tfrac{F}{\mathbb{P}^4_{[3]5}}]\!]\,\mathbb{P}^4_{[3]5} + [\![\tfrac{F}{\mathbb{P}^6_{[3]45}}]\!]\,\mathbb{P}^6_{[3]45} + \cdots \\
&= \overline{F} + [\![\tfrac{F}{\mathbb{P}^3}]\!]\,\mathbb{P}^3 + [\![\tfrac{F}{\mathbb{P}^1_3}]\!]\,\mathbb{P}^1_3 + [\![\tfrac{F}{\mathbb{P}^2_{13}}]\!]\,\mathbb{P}^2_{13} + [\![\tfrac{F}{\mathbb{P}^6_{[3]}}]\!]\,\mathbb{P}^6_{[3]} + [\![\tfrac{F}{\mathbb{P}^4_{[3]6}}]\!]\,\mathbb{P}^4_{[3]6} + [\![\tfrac{F}{\mathbb{P}^5_{[3]46}}]\!]\,\mathbb{P}^5_{[3]46} + \cdots
\end{aligned} \tag{15}
$$

Here, $\mathbb{P}^k_{[3]}$ means a polynomial in k that is orthogonal to traits $\phi_1$ through $\phi_3$. The blue terms are for the traits describing site 1, and the yellow terms are for the traits describing site 2. Since all of the terms for site 2 are independent of all of the terms for site one, we could combine any of the three sets of yellow terms with any of the sets of blue terms.

The fact that we can write the result in terms of different sequences is a consequence of the fact that there are multiple ways to construct orthogonal polynomials for these traits. Within a site, we could start with any of the three traits, then construct polynomials for the others that are orthogonal to it. *The key to our approach is to recognize that one of these sets of polynomials for a site is preferable to the others.*

There is a distinct pattern of correlation between the three traits associated with a single site (such as $\phi_1$, $\phi_2$, and $\phi_3$), since one of them must have a value of 1 and the others 0. This means that, when we construct orthogonal polynomials within a site, if we start with the "trait" corresponding to the nucleotide that is actually there, we know that the other "traits" at that site must have a value of zero. For example, if the nucleotide at site 1 is a $B$, then $\mathbb{P}^2$ is nonzero, but $\mathbb{P}^1_2$ and $\mathbb{P}^3_{12}$ are both zero, since the values of $\phi_1$ and $\phi_3$ are completely specified once we say that $\phi_2 = 1$. (I show this formally in the next section).

This means that, if we choose the right set of polynomials for a particular site, we need only consider one term in that set, since the others are zero. This preferred set of polynomials for a site is the one that starts with the nucleotide that is actually at that site, then calculates the values for the other possible nucleotides independent of that one. If our sequence is $BC$, this means that we can cancel some terms from the list of equations for $F$:

$$
F = \overline{F} + \left\{ \begin{array}{c}
[\![\tfrac{F}{\mathbb{P}^1}]\!]\,\mathbb{P}^1 + [\![\tfrac{F}{\mathbb{P}^2_1}]\!]\,\mathbb{P}^2_1 + [\![\tfrac{F}{\mathbb{P}^3_{12}}]\!]\,\mathbb{P}^3_{12} \\
\text{or} \\
[\![\tfrac{F}{\mathbb{P}^2}]\!]\,\mathbb{P}^2 + \cancelto{0}{[\![\tfrac{F}{\mathbb{P}^1_2}]\!]\,\mathbb{P}^1_2} + \cancelto{0}{[\![\tfrac{F}{\mathbb{P}^3_{12}}]\!]\,\mathbb{P}^3_{12}} \\
\text{or} \\
[\![\tfrac{F}{\mathbb{P}^3}]\!]\,\mathbb{P}^3 + [\![\tfrac{F}{\mathbb{P}^1_3}]\!]\,\mathbb{P}^1_3 + [\![\tfrac{F}{\mathbb{P}^2_{13}}]\!]\,\mathbb{P}^2_{13}
\end{array} \right. + \left\{ \begin{array}{c}
[\![\tfrac{F}{\mathbb{P}^4_{[3]}}]\!]\,\mathbb{P}^4_{[3]} + [\![\tfrac{F}{\mathbb{P}^5_{[3]4}}]\!]\,\mathbb{P}^5_{[3]4} + [\![\tfrac{F}{\mathbb{P}^6_{[3]45}}]\!]\,\mathbb{P}^6_{[3]45} \\
\text{or} \\
[\![\tfrac{F}{\mathbb{P}^5_{[3]}}]\!]\,\mathbb{P}^5_{[3]} + [\![\tfrac{F}{\mathbb{P}^4_{[3]5}}]\!]\,\mathbb{P}^4_{[3]5} + [\![\tfrac{F}{\mathbb{P}^6_{[3]45}}]\!]\,\mathbb{P}^6_{[3]45} \\
\text{or} \\
[\![\tfrac{F}{\mathbb{P}^6_{[3]}}]\!]\,\mathbb{P}^6_{[3]} + \cancelto{0}{[\![\tfrac{F}{\mathbb{P}^4_{[3]6}}]\!]\,\mathbb{P}^4_{[3]6}} + \cancelto{0}{[\![\tfrac{F}{\mathbb{P}^5_{[3]46}}]\!]\,\mathbb{P}^5_{[3]46}}
\end{array} \right. + \cdots \tag{16}
$$

Since we can choose any of the sums of blue terms, and combine this with any of the sums of yellow terms, we can write, for the sequence $BC$:

$$
F = \overline{F} + [\![\tfrac{F}{\mathbb{P}^2}]\!]\,\mathbb{P}^2 + [\![\tfrac{F}{\mathbb{P}^6_{[3]}}]\!]\,\mathbb{P}^6_{[3]} + 2^{nd}\,\text{order terms} \tag{17}
$$

Thus, our equation becomes much simpler if we can use the identity of the nucleotide that is present to pick out the right set of orthogonal terms. This is the function of the $\mu \otimes \mu$ terms in the main discussion; they pick out the nucleotide that is actually present at the site, essentially choosing the "preferred" set of terms for that site.

#### Proof that $\mathbb{P}_1^2 = 0$ when $\phi_1 = 1$

The approach discussed above hinges on the method of constructing an orthogonal basis within a site (by which we mean: over the set of possible nucleotides at that site). We need to prove that, if we start with the nucleotide that is actually at the site, the orthogonal polynomials corresponding to all other possible nucleotides are '0'.

Assume that the nucleotide at the site in question is $A$, so $\phi_1 = 1$. We construct the polynomial for $B$ independent of $A$ as:

$$\mathbb{P}_1^2 = \mathbb{P}^2 - \left[\!\left[\begin{smallmatrix}\mathbb{P}^2\\ \mathbb{P}^1\end{smallmatrix}\right]\!\right] \mathbb{P}^1 \tag{18}$$

Since we stipulated that $\phi_1 = 1$, we know that $\phi_2 = \phi_3 = 0$. From which it follows from Equation 14 that:

$$\begin{aligned} \mathbb{P}^1 &= 1 - \overline{\phi_1} \\ \mathbb{P}^2 &= -\overline{\phi_2} \end{aligned} \tag{19}$$

Substituting Equations 19 into Equation 18 yields:

$$\mathbb{P}_1^2 = -\overline{\phi_2} - \left[\!\left[\begin{smallmatrix}\mathbb{P}^2\\ \mathbb{P}^1\end{smallmatrix}\right]\!\right] (1 - \overline{\phi_1}) \tag{20}$$

We now need to evaluate the regression in Equation 20; which, by definition, is:

$$\left[\!\left[\begin{smallmatrix}\mathbb{P}^2\\ \mathbb{P}^1\end{smallmatrix}\right]\!\right] = \frac{[\![\mathbb{P}^1, \mathbb{P}^2]\!]}{[\![{}^2\mathbb{P}_1]\!]} \tag{21}$$

Since the $\mathbb{P}^i$ terms differ from the corresponding $\phi_i$ terms by a constant (Equation 14):

$$[\![\mathbb{P}^1, \mathbb{P}^2]\!] = [\![\phi_1, \phi_2]\!] = \overline{\phi_1\phi_2} - \overline{\phi_1} \cdot \overline{\phi_2} \tag{22}$$

For any site, at least one of $\phi_1$ or $\phi_2$ must be equal to 0 (as there can be only one nucleotide at any one site), so it must be the case that $\phi_1\phi_2 = 0$ for all sites for all individuals. We thus conclude:

$$[\![\mathbb{P}^1, \mathbb{P}^2]\!] = -\overline{\phi_1} \cdot \overline{\phi_2} \tag{23}$$

We now need the variance in $\mathbb{P}^1$:

$$[\![{}^2\mathbb{P}^1]\!] = \overline{\phi_1^2} - \overline{\phi_1}^2 \tag{24}$$

Since $\phi_i$ is either 0 or 1, for all $i$, it follows that $\phi_i^2 = \phi_i$ for each site for each individual. We can now rewrite Equation 24 as:

$$[\![{}^2\mathbb{P}^1]\!] = \overline{\phi_1} - \overline{\phi_1}^2 = \overline{\phi_1}(1 - \overline{\phi_1}) \tag{25}$$

Substituting Equations 23 and 25 into Equation 21 gives us the equation for the regression:

$$\left[\!\!\left[\begin{smallmatrix}\mathbb{P}^2\\ \mathbb{P}^1\end{smallmatrix}\right]\!\!\right] = -\frac{\overline{\phi_2}}{1 - \overline{\phi_1}} \tag{26}$$

Substituting Equation 26 back into Equation 20 now yields:

$$\mathbb{P}_1^2 = -\overline{\phi_2} + \frac{\overline{\phi_2}}{1 - \overline{\phi_1}}(1 - \overline{\phi_1}) = 0 \tag{27}$$

The polynomial for C independent of A and B is just:

$$\mathbb{P}_{1,2}^3 = \mathbb{P}^3 - \left[\!\!\left[\begin{smallmatrix}\mathbb{P}^3\\ \mathbb{P}_1^2\end{smallmatrix}\right]\!\!\right] \mathbb{P}_1^2 - \left[\!\!\left[\begin{smallmatrix}\mathbb{P}^3\\ \mathbb{P}^1\end{smallmatrix}\right]\!\!\right] \mathbb{P}^1 \tag{28}$$

Given the result in Equation 27, the same reasoning as above yields $\mathbb{P}_{1,2}^3 = 0$, and similarly for all possible monomers that are not present at the site.
